# Multicellular magnetotactic bacterial consortia are metabolically differentiated and not clonal

**DOI:** 10.1101/2023.11.27.568837

**Authors:** George A. Schaible, Zackary J. Jay, John Cliff, Frederik Schulz, Colin Gauvin, Danielle Goudeau, Rex R. Malmstrom, S. Emil Ruff, Virginia Edgcomb, Roland Hatzenpichler

## Abstract

Consortia of multicellular magnetotactic bacteria (MMB) are currently the only known example of bacteria without a unicellular stage in their life cycle. Because of their recalcitrance to cultivation, most previous studies of MMB have been limited to microscopic observations. To study the biology of these unique organisms in more detail, we use multiple culture-independent approaches to analyze the genomics and physiology of MMB consortia at single cell resolution. We separately sequenced the metagenomes of 22 individual MMB consortia, representing eight new species, and quantified the genetic diversity within each MMB consortium. This revealed that, counter to conventional views, cells within MMB consortia are not clonal. Single consortia metagenomes were then used to reconstruct the species-specific metabolic potential and infer the physiological capabilities of MMB. To validate genomic predictions, we performed stable isotope probing (SIP) experiments and interrogated MMB consortia using fluorescence *in situ* hybridization (FISH) combined with nano-scale secondary ion mass spectrometry (NanoSIMS). By coupling FISH with bioorthogonal non-canonical amino acid tagging (BONCAT) we explored their *in situ* activity as well as variation of protein synthesis within cells. We demonstrate that MMB consortia are mixotrophic sulfate reducers and that they exhibit metabolic differentiation between individual cells, suggesting that MMB consortia are more complex than previously thought. These findings expand our understanding of MMB diversity, ecology, genomics, and physiology, as well as offer insights into the mechanisms underpinning the multicellular nature of their unique lifestyle.

**Significance statement:** The emergence of multicellular lifeforms represents a pivotal milestone in Earth’s history, ushering in a new era of biological complexity. Because of the relative scarcity of multicellularity in the domains *Bacteria* and *Archaea*, research on the evolution of multicellularity has predominantly focused on eukaryotic model organisms. In this study, we explored the complexity of the only known bacteria without a unicellular stage in their life cycle, consortia of multicellular magnetotactic bacteria (MMB). Genomic and physiological analyses revealed that cells within individual MMB consortia are not clonal and exhibit metabolic differentiation. This implies a higher level of complexity than previously assumed for MMB consortia, prompting a reevaluation of the evolutionary factors that have led to the emergence of multicellularity. Because of their unique biology MMB consortia are ideally suited to become a model system to explore the underpinnings of bacterial multicellularity.

## Introduction

Multicellular lifeforms are defined as organisms that are built from several or many cells of the same species (1, 2). Beyond this, other characteristics of multicellularity include a specific shape and organization, a lack of individual cell autonomy or competition between cells, and a display of cell-to-cell signaling and coordinated response to external stimuli (3). The transition from a single cell to a cooperative multicellular organism is an important evolutionary event that has independently occurred at least 25 times across the tree of life (2). This suggests that the development of multicellularity can occur in any species given proper selective pressure (4, 5). Prior research on the transition of unicellular to multicellular organisms has largely focused on eukaryotic model systems such as choanoflagellates (6), fungi (7), and algae (8). Multicellularity within the domain Bacteria is comparatively rare (9), yet this lifestyle likely first evolved approximately 2.5 billion years ago (10). Examples of multicellularity within the domain Bacteria include filamentous cyanobacteria (*e.g*., *Anabaena cylindrica*), mycelia-forming actinomyces (*e.g., Streptomyces coelicolor*), swarming myxobacteria (*e.g.*, *Myxococcus xanthus*), centimeter-long cable bacteria (*e.g*., *Electrothrix* sp.), and the recently discovered liquid-crystal colonies of *Neisseriaceae* (*e.g., Jeongeupia sacculi* sp. nov. HS-3) (5, 11, 12). While capable of multicellular growth, each of these microbes undergoes a unicellular stage at some point in their life cycle.

Currently, the only known example of purportedly obligate multicellularity – an organism without a detectable unicellular stage – within the domain Bacteria are several species of multicellular magnetotactic bacteria (MMB; we use the terms ‘MMB consortia’ and ‘MMB’ interchangeably) (13, 14). MMB are symmetrical single-species consortia composed of 15-86 cells (15)of Desulfobacterota (formerly Deltaproteobacteria) arranged in a single layer enveloping an acellular, central compartment (Fig. 1A-B). Consortia range in size from 3-12 µm in diameter (16–18). Within the Desulfobacterota, MMB form an uncultured, monophyletic family that is distinct from several physiologically and genetically well-characterized unicellular relatives, suggesting a common ancestor that achieved a multicellular state (19–21). MMB are globally distributed in sulfidic brackish and marine sediments but typically are of low relative abundance in these habitats (0.001 - 2% (18, 22, 23)). In addition to their unique obligate multicellular lifecycle, MMB have an organelle called the magnetosome (24). The magnetosome is a lipid vesicle that encapsulates biomineralized magnetite (Fe3O4) and/or greigite (Fe3S4, Fig. 1C) and allows MMB to sense and orient themselves along Earth’s geomagnetic field in a phenomenon termed magnetotaxis. Magnetosome formation is controlled by a magnetosome gene cluster (MGC, SI Appendix Text) that encodes several proteins involved in the formation, alignment, and maturation of the organelle (25, 26). The presence of magnetosomes in MMB can be exploited to physically enrich them from environmental samples using a magnet (SI Videos S1 and S2). This is particularly important considering that MMB have not yet been successfully cultured MMB are distinctive among bacteria because their life cycle lacks a unicellular stage. Instead, MMB replicate by the entire consortium doubling its cell number and volume before separating into two, seemingly identical consortia (14, 16, 27, 28). Historically, MMB have been described as “aggregates” of cells (29), which could imply that individual cells assemble to form a multicellular aggregate, akin to the early stages of biofilm formation (5, 29). In this study we use the terms “consortium” (singular) and “consortia” (plural) to describe the unique form of multicellularity observed for MMB.

**Fig. 1.**
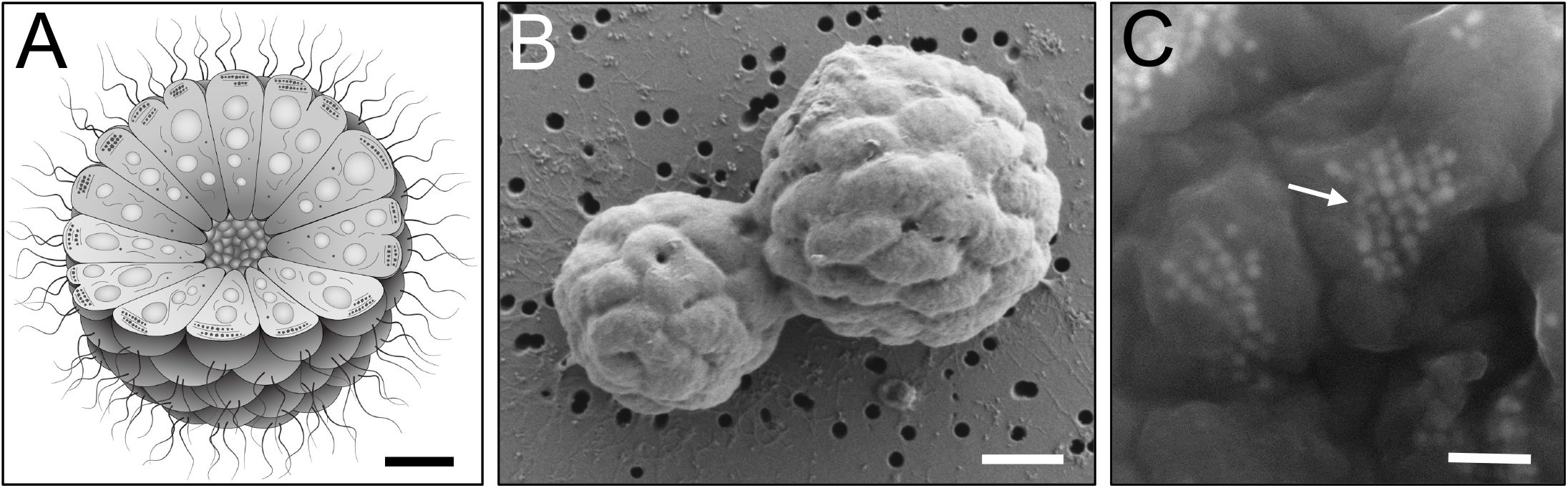
Morphology and structure of MMB. *(A)* Cartoon depicting the morphology and internal organization of a MMB consortium. At the center of each MMB consortium lies an acellular space that is surrounded by a single layer of cells. Each cell harbors magnetosome organelles (black polygons aligned along cytoskeleton-like filaments), compartments for carbon or energy storage (gray circles), as well as other, currently unidentified structures. Scale bar ca. 1 µm. *(B)* Scanning electron microscopy (SEM) image of two MMB magnetically enriched from LSSM, possibly undergoing division. Scale bar, 1 µm. *(C)* Backscatter electron microscopy image of magnetosome chains within MMB cells (arrow). Magnetosome minerals appear to have 4-8 visible facets and are approximately 30-60 nm in diameter. Scale bar, 300 nm. Contrast and brightness of image *(C)* was increased for better visualization.

Under external stress, an MMB consortium becomes dismantled, followed by an immediate loss of magnetic orientation and motility and eventual loss of membrane integrity, leading to cell death (30). MMB consortia consistently exhibit a high degree of magnetic optimization, excluding the possibility that the consortium is a mere aggregation of cells without underlying self-organization (31, 32). Each cell within the consortium has multiple flagella, resulting in the whole consortium being peritrichously flagellated (17, 33). When environmental conditions change, such as alterations in light exposure or magnetic fields, a coordinated response in motility occurs within fractions of a second (33, 34). This collective response implies inter-cellular communication among individual cells, which is hypothesized to occur through the central acellular volume that the cells surround (16). Previous work has hypothesized that the absence of a single cell stage in MMB might be necessary to maintain the acellular volume at the center of each MMB or that their larger size is needed to evade predation by protists (14). Currently, there is no evidence to support or refute these hypotheses. While past studies have presented fascinating insights into the cellular organization of MMB and their diverse abilities to sense the environment via light and electron microscopy (20, 34, 35), their recalcitrance to cultivation has hindered progress towards a better understanding of their physiology and genomics. With the exception of a study that demonstrated chemotactic response of MMB consortia to small molecular weight organic acids (35), questions about their physiology remain unaddressed, and hypotheses about the potential for metabolic differentiation or a division of labor between individual cells within a consortium have not been experimentally tested.

To address these knowledge gaps, we investigated the taxonomic diversity, genomics, physiology, metabolic differentiation, and clonality of MMB inhabiting a tidal pool. To investigate the diversity of MMB within this environment, we sequenced the Single Consortium Metagenomes (SCMs) of 22 MMB consortia, representing eight distinct species of MMB. Comparing the SCMs we were able to quantify the extent of single nucleotide polymorphisms (SNPs) between cells composing individual MMB consortia. Our analyses showed that MMB exhibit genetic diversity within a single consortium, indicating that they are not composed of clonal cells. Physiological predictions were established through the reconstruction of species-specific metabolic models. We tested these predictions by performing stable isotope probing (SIP) experiments and analyzing individual consortia using fluorescence *in situ* hybridization (FISH), nano-scale secondary ion mass spectrometry (NanoSIMS), and bioorthogonal non-canonical amino acid tagging (BONCAT). Our results demonstrate that MMB are mixotrophic sulfate reducers and that individual cells within MMB consortia exhibit dramatically different rates of substrate uptake, indicating metabolic differentiation, as well as localized protein synthesis activity.

## Results and Discussion

### Genomic features and phylogenetic analysis of MMB

MMB were recovered from sulfidic sediments collected from a tidal pool in Little Sippewissett Salt Marsh (LSSM; Falmouth, MA, Fig. S1A-B). This sample site was selected based on the ability to magnetically enrich (SI Videos S1 and S2) relatively large quantities of MMB, as previously demonstrated (34, 36). Individual MMB consortia were sorted from a magnetically enriched pellet using fluorescence-activated cell sorting and the DNA of individual sorted MMB was amplified by multiple displacement amplification before Illumina sequencing. From this sample, the SCMs of 22 individual MMB were recovered (Fig. 2, SI Appendix Table S1). The GC content of the SCMs ranged from 36.2 to 38.4%, which is similar to the GC content observed in previously published MMB draft genomes (20, 37, 38). The average and median size of the 22 new SCMs was 7.7 Mb, with a range from 6.1 to 9.1 Mb (SI Appendix Table S1). Prior to this study, only three draft genomes of MMB had been sequenced. These genomes exhibited significant variations in size, ranging from 14.3 Mb for *Ca*. Magnetomorum sp. HK-1 (37), 12.5 Mb for *Ca*. Magnetoglobus multicellularis (20), and 8.5 Mb for MMP XL-1 (38), although the MMP XL-1 genome is not publicly available. The genome sizes of *Ca*. M. multicellularis and *Ca*. M. sp. HK- 1 could be conflated due to contamination or the combination of sequence data into the same final bin, as discussed in the respective studies (20, 37) and evidenced by our own evaluations of genome contamination (Fig. 2A) Only 14 of the 22 SCMs contained 16S rRNA genes (SI Appendix Table S1). These sequences, together with publicly available 16S rRNA sequences of MMB as well as those of their single-cell relatives *Desulfosarcina variabilis* and *Ca*. Desulfamplus magnetomortis BW-1, were used to construct a phylogenetic tree (SI Appendix Table S2). This analysis revealed the presence of five phylogenetically distinct genera of MMB in LSSM with high bootstrap support (>75%) (Fig. S2). Analysis of amplicon sequence data obtained in this study and sequences from a previous study at LSSM (36) showed that Group 1 MMB was most abundant in the sample site, constituting 61% of all 16S rRNA genes. Groups 2, 4, 5, and 3 accounted for 21%, 6.5%, 6.5%, and 5% of the 16S rRNA genes, respectively (Fig. S2, S3).

**Fig. 2.**
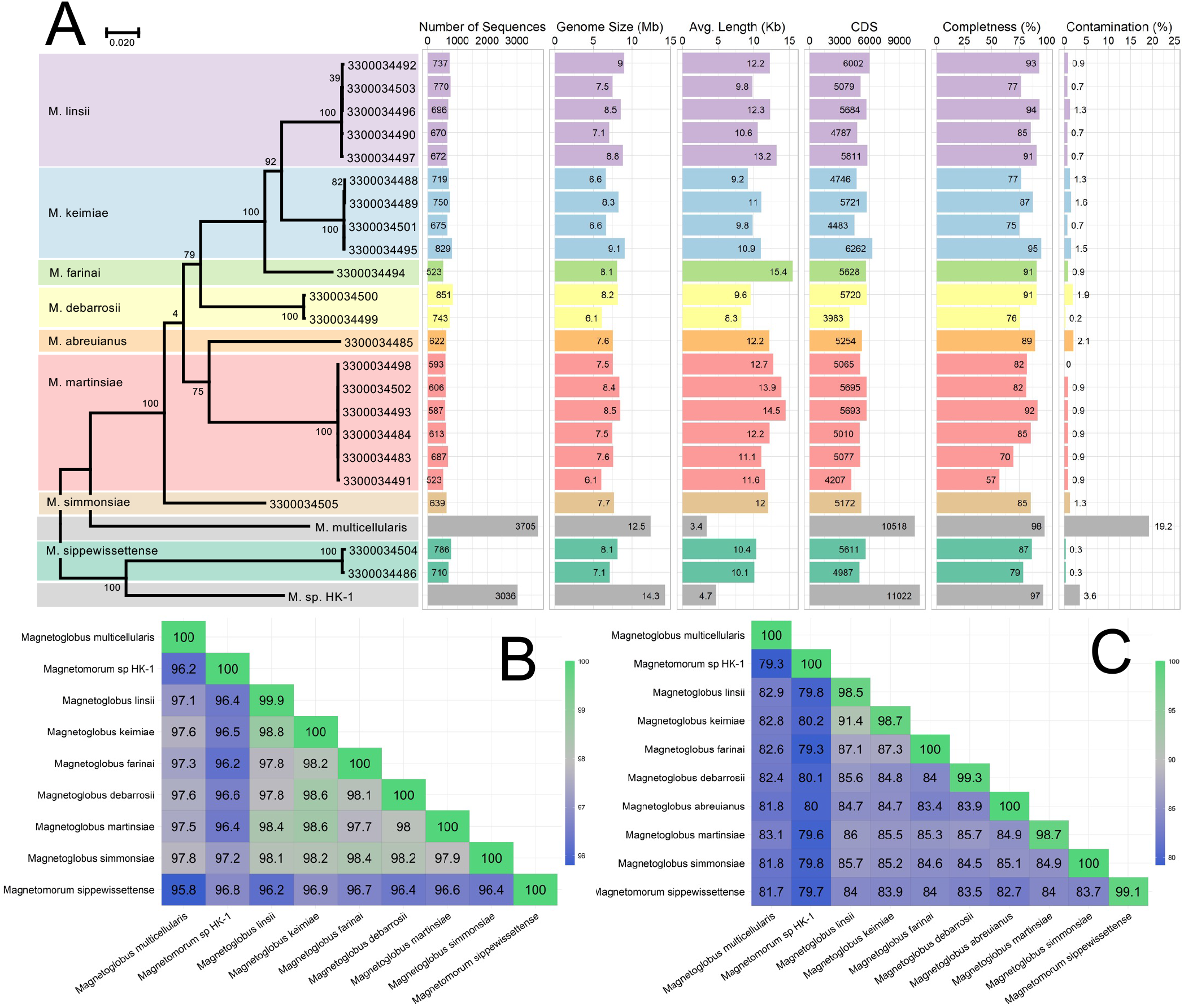
Genomic and phylogenetic analysis of all publicly available MMB MAGs and the 22 SCMs generated in this study. *(A)* Maximum-likelihood tree, inferred with FastTree, using a concatenated set of six conserved COGs (Table S3) present in all entries. Ultrafast bootstrap support values and selected genome statistics are listed. The color codes for the SCM Groups remain the same throughout all figures. *(B)* Average full length 16S rRNA gene identity and *(C)* average genome nucleotide identity heat maps of the eight newly identified MMB species compared to two available MMB reference genomes (*Ca.* M. multicellularis and *Ca.* Magnetomorum sp. HK-1). For a phylogenetic tree of all publicly available MMB 16S rRNA gene sequences, see Fig. S2. For an exhaustive sequence identity analyses of 16S rRNA and whole genomes of MMB see Figs. S3-5.

Phylogenomic analysis of six bacterial single copy genes found in all recovered MMB SCMs yielded a topology consistent with the phylogeny derived from the 16S rRNA gene sequences (Fig. 2, SI Appendix Table S3, Fig. S2). Similarly, whole genome and 16S rRNA specific ANI analyses resolved eight unique species of MMB with >96% average nucleotide identity. We assigned type genomes for each new MMB species and named them after scientists who have greatly advanced our knowledge of MMB (SI Appendix Text, SI Appendix Table S4).

### Clonality within MMB

MMB have historically been assumed to be clonal due to the synchronized replication of cells during division, which should result in genetically identical daughter cells in the same consortium (14, 27). Additionally, obligate multicellularity has traditionally been thought to perpetuate a clonal population (39). Although MMB maintain an obligate multicellular lifecycle, the degree to which clonality exists within a single consortium has never been experimentally tested. Currently, the only evidence suggesting that cells within MMB are closely related comes from analyses of the 16S rRNA genes from cells of a single genome amplified MMB consortium (37) and a FISH study demonstrating that cells within individual MMB have identical 16S rRNA sequences (36).

We set out to test the hypothesis of clonality using comparative genomics of the 22 MMB SCMs recovered in this study. Reads from each individual SCM were mapped to the corresponding genome bins to quantify single nucleotide polymorphisms (SNPs) within a single MMB consortium. As a procedural control, 10, 30, 60, and 100 cells of a clonal culture of *Pseudomonas putida* were sorted to construct a mock multicellular consortium. The DNA of MMB consortia and *P. putida* controls were amplified using multiple displacement amplification and sequenced using Illumina short read sequencing. Our analysis of the SCMs revealed for the first time that MMB consortia are genomically heterogeneous and thus do not fit the model of clonality for obligate multicellular organisms (Fig. 3A). MMB from LSSM contain up to two orders of magnitude more SNP differences within a single consortium as compared to the same number of cells from the clonal control (p < 7.3 x 10^-9^), with an estimated range of 157-789 SNPs in individual SCMs (Fig. 3, SI Appendix Table S5). Other environmental microbes co-sorted with MMB showed a SNP rate similar to the clonal control and a SNP rate statistically different from the MMB (p < 2.4 x 10^-6^), illustrating the uniqueness of MMB. Wielgoss *et al.* performed a similar analysis on fruiting bodies of the aggregative multicellular bacterium *Myxococcus xanthus* in which a comparison of the genomes of cells in fruiting bodies revealed 30 SNP differences between lineages originated from a recent single ancestral genotype (40). Furthermore, nearly half the mutations detected in the *M. xanthus* genomes occurred in the same six genes, suggesting there was a strong selection for socially relevant genes, such as a histidine kinase (signal transduction) and methyltransferase (gene expression). Positive selection upon cooperative genes may promote diversity within the organism as a mechanism to increase fitness within spatiotemporally variable environments and protect against social cheaters (41).

**Fig. 3.**
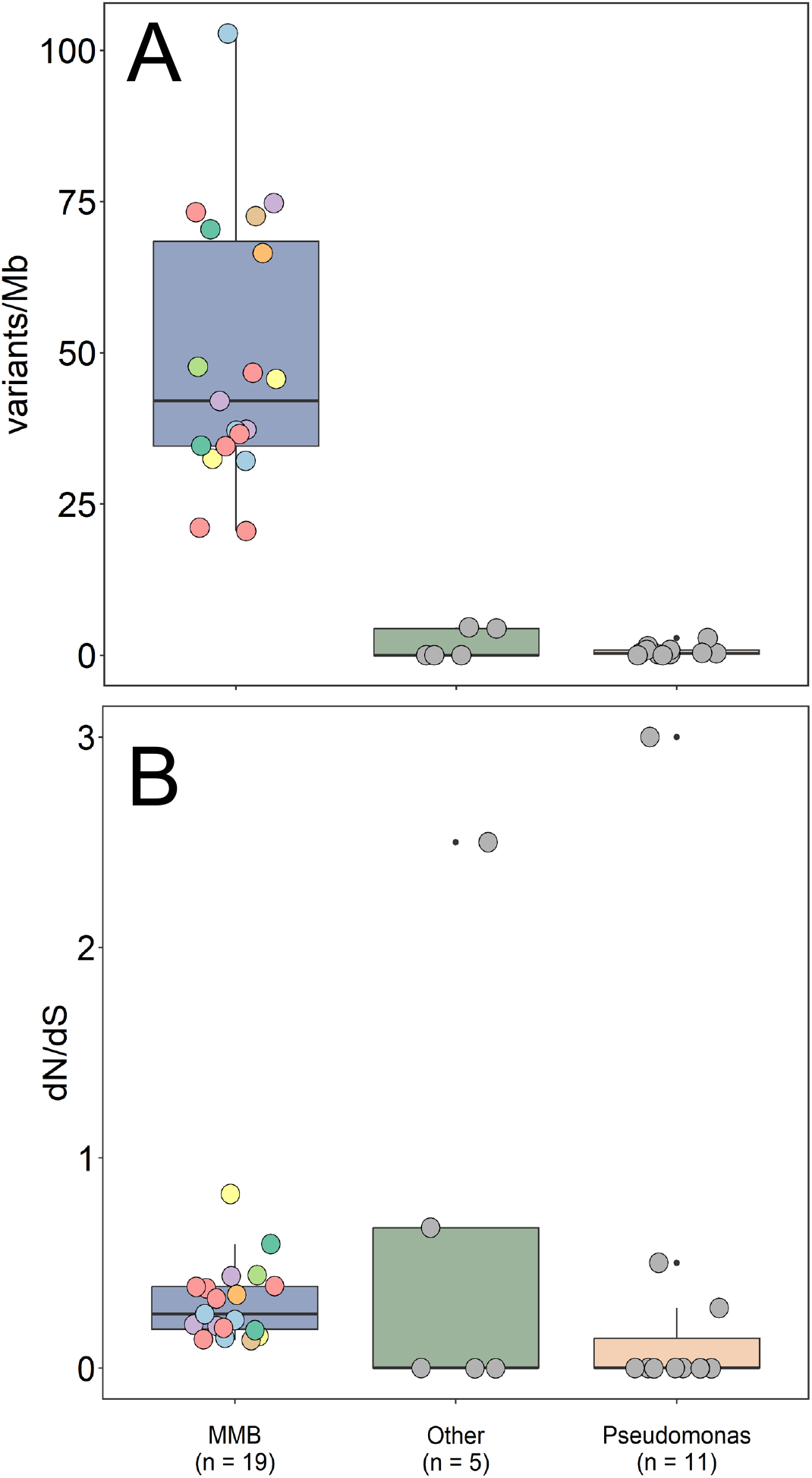
Clonality analysis of individual MMB consortia. (*A*) Individual reads were mapped to the same genome bin for each of the 22 SCMs. This analysis revealed that the genomes of cells within MMB consortia have a higher single nucleotide polymorphism rate (SNP expressed as Variations per kb) as compared to a clonal *Pseudomonas* sp. control (p < 7.3 x 10^-9^, n = 10, 30, 60, and 100 *Pseudomonas* cells) and other environmental cells (p < 2.4 x 10^-6^, *e.g*. “Other”). (*B*) The three sample categories showed no statistically significant difference in terms of their ratio of non-synonymous to synonymous substitutions (dN/dS). Values near 0 indicate that substitutions are neutral and there is no positive selection of the protein-coding genes in which the SNPs reside. The color of each SCM corresponds to the color identifying each unique species in Fig. 2.

To investigate if the genetic heterogeneity within MMB contributes to an increased fitness of the organism, we identified the genes containing SNPs and calculated the corresponding ratio of non-synonymous (dN) to synonymous (dS) substitutions. This analysis showed that the SNP differences within the SCMs of MMB appear to be random with no single gene or category of genes exclusively impacted by the SNPs within or across MMB consortia (Fig. 3B, SI Appendix Table S6). SNPs with a high dN/dS ratio were predominantly found in unannotated genes, such as hypothetical proteins (Fig. S6). Such unannotated genes that are subject to stronger positive selection could ultimately drive functional divergence within the consortium. Other benefits of genomic heterogeneity within MMB are not readily apparent and could be attributed to errors during DNA replication or damaging effects of mutagens. However, it has been shown that a single mutation can lead to a division of labor in bacteria (42). At this point, it is unclear whether any of the changes we observe in the genomes contained within individual MMB would lead to phenotypic differentiation between the adjacent cells.

### Genome annotation

Metabolic reconstructions of the MMB SCMs (Fig. 4, SI Appendix Table S7) revealed that all MMB are capable of heterotrophic sulfate reduction and can use acetate, succinate, and propionate as carbon donors and/or electron sources, consistent with previous genomic analyses (20, 37). The SCMs show that LSSM MMB have highly similar metabolic potential. One exception is *Ca*. M. sippewissettense, which lacks the ability to utilize acetyl-coenzyme A (CoA) synthetase and is unable to use acetate, instead likely relying on lactate dehydrogenase to metabolize lactate, a substrate the other species are not capable of using. None of the SCMs contain acetaldehyde dehydrogenase, indicating that MMB are not capable of alcohol fermentation. We resolved a complete glycolysis pathway and TCA cycle as well as reductive CoA pathway in all SCMs. The presence of these genes suggests that MMB in LSSM are capable of both heterotrophic and autotrophic growth using sulfate reduction coupled to hydrogen metabolism, by means of *hyaA/B* and *hybA/B* complexes and oxidative phosphorylation. MMB are genetically capable of shuttling electrons using complexes I, II, and V of the oxidative phosphorylation pathway using F-type ATP synthase complexes, although partial V/A type ATP synthase were found in *Ca*. Magnetoglobus martinsiae and *Ca*. Magnetomorum sippewissettense. In addition, they encode a full Nqr (Na^+^- transporting NADH:ubiquinone oxidoreductase) complex that can move electrons from NADH to ubiquinone with the translocation of a Na^+^ across the membrane. Cytochrome bd oxidase subunits I and II are present in all SCMs, except *Ca*. Magnetoglobus farina, and could be used to respire molecular oxygen (O2) using electrons from cytochrome c or quinols (43). All species of MMB from LSSM encode rubrerythrin and superoxide reductase, suggesting the possibility that O2 could instead be detoxified by the cytochrome bd oxidase (SI Appendix Table S7) (20, 44). Electrons can also be removed by the reduction of protons to molecular hydrogen (H2) by group 1 nickel-iron hydrogenases. The H2 can then diffuse across the membrane where HybA/B could oxidize the H2, yielding two electrons and two protons. From there, cytochrome c can shuttle the electrons to the Dsr and Qmo complexes for dissimilatory sulfate reduction.

**Fig. 4.**
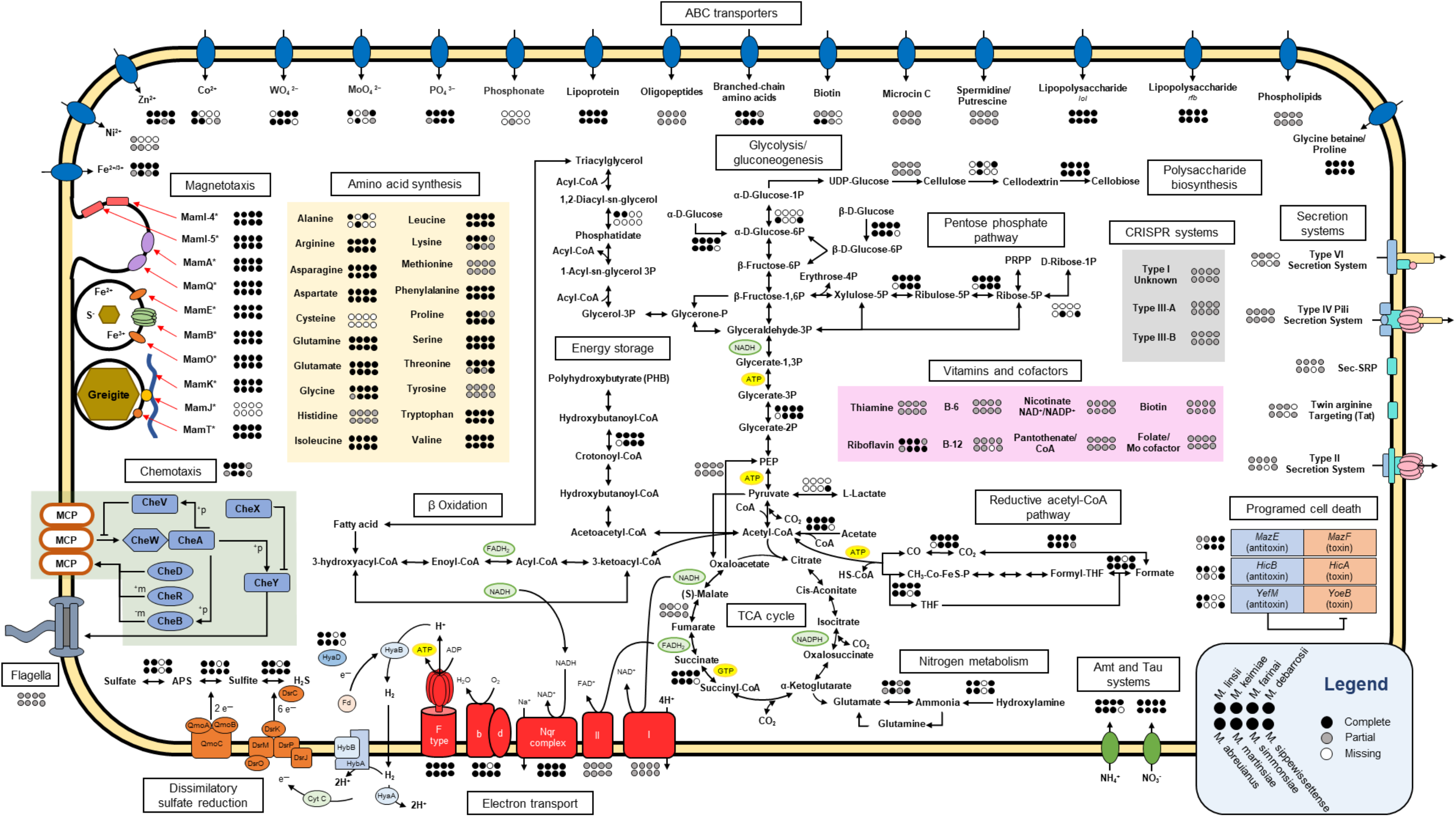
Metabolic potential of the eight MMB species in LSSM. Arrows without circles indicate presence of the respective enzyme or pathway in all bins. Circles indicate complete presence (black), partial presence (gray), or missing (white) genes in each species. A full list of genes used to construct this figure can be found in Table S5.

The MMB SCMs encode several divalent metal transporters, including FoaAB ferrous iron and FepBDC ferric iron transport proteins, indicating they are capable of using both Fe(II) and Fe(III). All SCMs encode phosphate transporters as well as oligopeptide and branched-chain amino acid transporters. Genes for polyamine transport were recovered in the SCMs and may provide resistance to environmental stress such as osmotic pressure and reactive oxygen species (45). Additionally, each SCM encodes a glycine betaine transporter but does not encode a betaine reductase, indicating that MMB do not use glycine betaine as a nitrogen source but as an osmoprotectant (46). All MMB species in LSSM, except *Ca.* M. sippewissettense, encode an Amt transporter to transport ammonia into cells that can then be converted into glutamine or glutamate and fed into anabolic pathways. Additionally, each species encodes the NitT/TauT system for nitrate, sulfonate, and bicarbonate transport into cells. The SCMs showed that MMB are capable of synthesizing all canonical amino acids except cysteine and lack cysteine prototrophy genes. Cultures of single celled magnetotactic bacteria have been found to require the addition of cysteine for growth, suggesting that many magnetotactic bacteria, including MMB, cannot synthesize their own cysteine (47). The inability to synthesize a sulfurous amino acid is surprising given that most magnetotactic bacteria, including all known MMB, live in sulfur-rich environments.

Previous studies using transmission electron microscopy have found large vesicles within MMB cells that have been attributed to carbon/energy or phosphate storage (48). Metabolic analysis of the SCMs showed that acetyl-CoA could be condensed and polymerized to polyhydroxybutyrate (PHB) for storage. Furthermore, all necessary genes were identified for *β*- oxidation using triacylglycerol synthesized from the acylation of glycerol-3P with acyl-CoA (Fig. 4, SI Appendix Table S7). Using Raman microspectroscopy applied to individual MMB, we demonstrated the presence of PHB and lipids, along with Nile Red staining of carbon-rich droplets within cells (Fig. S7, SI Appendix Table S8). This is, to our knowledge, the first time carbon and energy storage compounds in MMB have been unambiguously identified. Carbon storage has been shown to support the multicellular reproductive life cycles in *Vibrio splendidus* through the specialization of cells during resource limitations (49), suggesting that MMB may utilize a similar mechanism to support their multicellular growth.

Altruistic behavior in biological systems is often favored when relatedness among species is high and the benefit is comparatively large compared to the cost, as has been observed in multicellular myxobacteria (41). The SCMs revealed that MMB encode *mazE/F*, *hicA/B*, and *yefM/yefB* type II toxin-antitoxin (TA) systems (Fig. 4, SI Appendix Table S7). TA systems represent an extreme example of altruism in multicellular systems, as individual cells that contribute to the organism by sacrificing themselves through death do not directly benefit from the organism’s multicellularity. But, selection favoring altruistic traits occurs due to the fitness benefits those traits impart on relatives (50). Detection of CRISPR (clustered regularly interspaced short palindromic repeats) systems I, III-A, and III-B (SI Appendix Table S7) suggest the TA systems could be used in response to viral infection (51). The evolution of altruistic cooperation in multicellular organisms has been proposed as a response to environmental stressors (50), indicating the presence of TA systems likely confer increased fitness for MMB in the environment.

### Cell-to-cell adhesion

One of the most intriguing features of MMB is their multicellular lifecycle. But how these bacteria maintain their multicellular shape is not entirely known. Previous genomic and microscopic analysis of MMB suggested that exopolysaccharides, adhesion molecules, and Type IV pili could be involved in cell-to-cell adhesion (20, 52). Extracellular matrices, specifically those composed of polysaccharides, have been shown to be important for the development and maintenance of bacterial multicellularity, resulting in several emergent properties that benefit the organism, including the reduction of maintenance energy for individual cells (53). *Myxobacteria sp.* and *Escherichia coli* have both been shown to use exopolysaccharides to maintain macroscopic biofilms, (7, 54). The SCMs recovered in this study encode genes for extracellular polysaccharide biosynthesis, including family-2 glycosyltransferases (GT2), which have been shown to secrete diverse polysaccharides such as cellulose, alginate, and poly-N-acetylglucosamine (55, 56). Specifically, the genes identified in the SCMs were homologous to GT2 Bcs proteins, a bacterial protein complex that synthesizes and secretes a *β*-1,4-glucose polymer (*e.g*., cellulose) during biofilm formation (SI Appendix Table S7) (57, 58). The LSSM MMB encode enzymes that catalyze the production of cellulose for biofilm formation (*bcsA*, *bcsQ*, *bcsZ*, *pilZ*, and *bglX*), but lack the co-organization of genes at a single locus as observed for other bacteria (57). Furthermore, the *bcsB* and *bcsC* subunits were not identified, but additional GT2 as well as *wza* genes that may be involved in the synthesis of exopolysaccharides were present (59). The catalytic activity of BcsA has been shown to be influenced by the concentration of cyclic dimeric guanosine monophosphate (c-di-GMP) which is in turn affected by environmental oxygen levels (60, 61). Under oxic conditions the cellular level of c-di-GMP has been shown to increase and bind to BcsA, leading to increased cellulose synthesis (61). Because MMB commonly exist in oxygen-deficient sediments, cellulose synthesis may be triggered under oxic conditions to stimulate biofilm formation, which has been observed in cultivation attempts of MMB (20).

Filamentous hemagglutinin has been shown to recognize and bind to carbohydrates to facilitate cell-to-cell adhesion in a biofilm (62, 63). The presence of filamentous hemagglutinin genes in our SCMs suggests MMB could use these protein complexes as a mechanism for cell-to-cell adhesion, as previously suggested (20). Furthermore, the SCMs encode genes for OmpA/F porins, proteins with adhesive properties that have been suggested to interact with exopolysaccharides leading to aggregation of cells (64). Type IV pili, which have been shown to be involved in cell-to-cell adhesion by interacting with exopolysaccharides (65), were also identified in the SCMs. The pili could alternatively be used for motility, chemotaxis, organization, and DNA uptake (66). Further investigation into the use of the Type IV pili within MMB is warranted as only predictions can be made from the available genomes.

Previous studies on the membrane of MMB using Ruthenium Red dye and calcium cytochemistry have shown that the consortia are coated in a polysaccharide that extends between cells into the acellular central compartment but the exact composition and structure of this polysaccharide remains unclear (16, 67). Using Raman microspectroscopy we identified peaks corresponding to exopolysaccharides, confirming the presence of an exopolysaccharide within or surrounding MMB (Confocal Raman does not have enough z-resolution to distinguish the in- and out-side of cells; Fig. S7, Appendix Table S8). Cellulase hydrolysis of the MMB resulted in eroded surfaces of the consortia, demonstrating that MMB are indeed covered by a cellulose layer (Fig. S8). Together, these analyses highlight the structural and functional significance of exopolysaccharides required for the multicellular morphotype of MMB.

### Abundance, distribution, and *in situ* activity of MMB in LSSM

Temporal shifts in MMB groups at LSSM have previously been documented (68) but the abundance of MMB correlated to sediment depth has not yet been analyzed. MMB in the LSSM subsurface were quantified by retrieving a 15 cm core from the tidal pond and determining the fractional abundance of each of the five MMB groups recovered throughout the core at centimeter-scale resolution using newly designed fluorescence *in situ* hybridization (FISH) probes (Fig. S9, SI Appendix Table S9). In the top five centimeters of sediment, Group 1 MMB accounted for >75% of all MMB while the other groups accounted for 1-25%, depending on sediment depth. The total abundance of MMB dropped sharply below 5 cm, where the sediment horizons transitioned from sandy to dense clay sediment containing plant roots. This could be due to MMBs preference for low oxygen conditions, under which sulfate reduction is favored (35, 69). A similar depth- abundance profile was previously observed for the closely related MMB *Ca*. M. multicellularis (69).

Bioorthogonal noncanonical amino acid tagging (BONCAT) was used to determine the anabolic activity of MMB Group 1 in the top 6 cm of the LSSM core, which hosted the majority of MMB. Using this approach, we identified a statistically significant difference in MMB activity from 1 cm depth compared to the 2-3 cm (p < 3.4x10^-4^) and from 3 cm compared to 4-5 cm (p < 3.9x10^-3^), below which the MMB population diminished (Fig. S10). The increase of activity of MMB in the first 5 cm of the sediment could be attributed to the circumneutral pH and low redox potential (-260 to -460 mV), as previously observed to be important for the bioavailability of iron and sulfur species for MMB (37).

### Physiology of MMB

Previous genome- and chemotaxis-based studies suggested that MMB live by heterotrophic sulfate reduction using small organic acids as electron donors (20, 35, 37). However, no direct observation of the use of such organics has been reported. Our metabolic reconstructions revealed that all MMB species in LSSM are genetically capable of coupling sulfate reduction to the use of acetate, propionate, and succinate as well as inorganic carbon fixation via the reductive acetyl-CoA pathway. To test whether MMB use these carbon sources to support their growth, we incubated sediment samples with ^13^C-labeled substrates (acetate, bicarbonate, propionate, and succinate) *in situ* and analyzed individual MMB using Nano-scale secondary ion mass spectrometry (NanoSIMS). MMB that had been incubated with ^13^C-acetate exhibited higher ^13^C labeling as compared to the other substrates, which could suggest a preference for acetate (Fig. 5, SI Appendix Table S10). To identify specific MMB groups, FISH was performed prior to NanoSIMS analyses. Group 1 MMB showed the highest incorporation of ^13^C from acetate as compared to Groups 3 and 4 (p < 1.5x10^-3^, Fig. S11). We also observed a significant difference between Group 1 and 4 for ^13^C-bicarbonate and ^13^C-propionate uptake (p < 3.9x10^-3^ and 5.8x10^-5^, respectively). At least three genera of MMB (*i.e.*, Groups 1, 2, and 3) assimilated both bicarbonate and propionate (Fig. S15). We were unable to magnetically enrich MMB from a sediment sample incubated with ^13^C-acetate and molybdate, an inhibitor of sulfate reduction, indirectly demonstrating that MMB are in fact sulfate reducers. In summary, our analyses demonstrated that LSSM MMB are capable of assimilating both inorganic and organic carbon, indicating autotrophic and heterotrophic growth, and that different Groups of MMB demonstrate variable affinities for carbon sources.

**Fig. 5.**
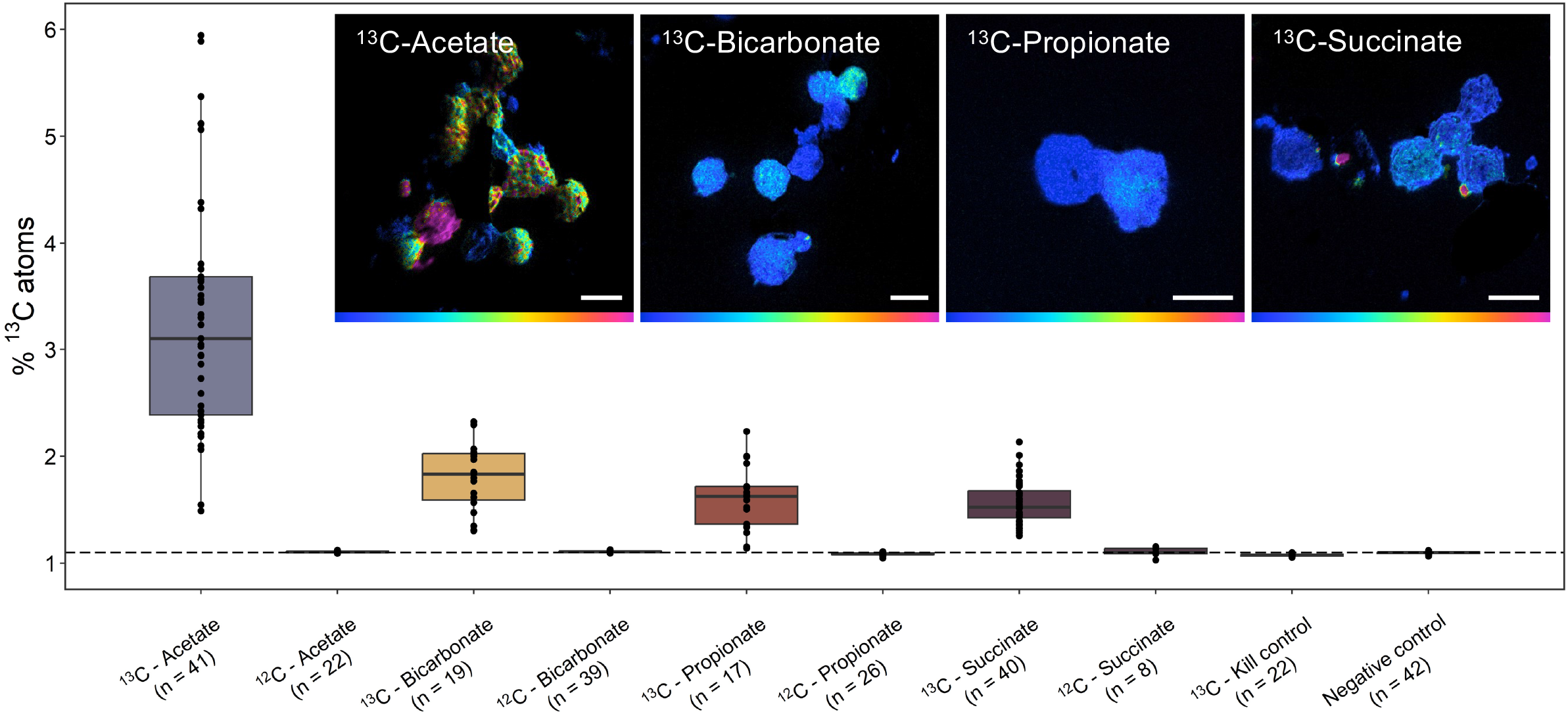
NanoSIMS analysis of the cellular ^13^C-content of MMB consortia after *in situ* incubation with isotopically light or heavy carbon sources, specifically 1,2-^13^C_2_-acetate, ^13^C-bicarbonate, 1,2-^13^C_2_- propionate, or 1,2-^13^C_2-_succinate, for 24 hours. The kill control contained magnetically enriched MMB that had been fixed in 4% paraformaldehyde prior to ^13^C-acetate addition. The negative control was sediment containing MMB without substrate addition. The dotted line shows the natural abundance of ^13^C. For further description of boxplots, see SI Appendix Text. Inset images show representative NanoSIMS hue saturated images (HSI) for each ^13^C-labeled substrate analyzed. Color scales in HSI images are 1.1% - 5% atom percent ^13^C. Scale bars are 5 μm. Fig. S1CD show the incubation setup. For a comparison of the anabolic activity of MMB groups 1, 3, and 4 see Fig. S11. Fig. S12 provides an example for correlative microscopy analysis of MMB. SI Materials and Methods detail the calculation of atom percent. For ROIs, see Fig. S13.

### Metabolic differentiation as studied by SIP-NanoSIMS

A hallmark of multicellularity is the existence of a division of labor (5), however, because of their recalcitrance to cultivation, this hypothesis has never been addressed in MMB. To investigate whether MMB are metabolically differentiated, a magnetic enrichment of MMB was incubated *in vitro* with ^13^C-labeled acetate and deuterium oxide (^2^H2O), with cellular labelling from the latter being a general proxy for metabolic activity (70). Samples analyzed using NanoSIMS showed variation of isotopic signal across cells within individual consortia, indicating different metabolic activity within MMB (Fig. 6, SI Appendix Table S11). The mass ratio for each isotope label was quantified and areas of high anabolism (referred to as “hotspots”) within the consortium compared to the value of the same isotope label for the whole consortium. This analysis demonstrated a statistically significant difference of anabolic activity between hotspots and whole consortium for both ^13^C and ^2^H2O (p < 1.3x10^-3^ and < 2.2x10^-8^, respectively). Comparison of SEM and NanoSIMS imaging shows that the extent of SIP labeling varies within a single cell as well as across the entire MMB consortium (Fig. S12). The hotspots do not exhibit localization in any specific region of an MMB. However, they are not uniformly distributed throughout the consortium, demonstrating variations in metabolic activity with some areas displaying lower metabolic activity than others. To further investigate the localization of the isotope within the individual consortium, we applied a median filter ratio to the hue saturated images (HSI) using different kernel radii (71). This method averages the isotopic ratio over the given pixel radius, revealing sub-consortium localization across the MMB (Fig. S15). Together, our analyses shows that metabolism of ^13^C-acetate and ^2^H-water is not uniform across the MMB, suggesting a differentiation in metabolic activity within individual consortia. Similar differences in the uptake of isotope-labeled substrate have also been reported for cellularly and metabolically differentiated cells of filamentous cyanobacterium *Anabaena oscillarioides* (72).

**Fig. 6.**
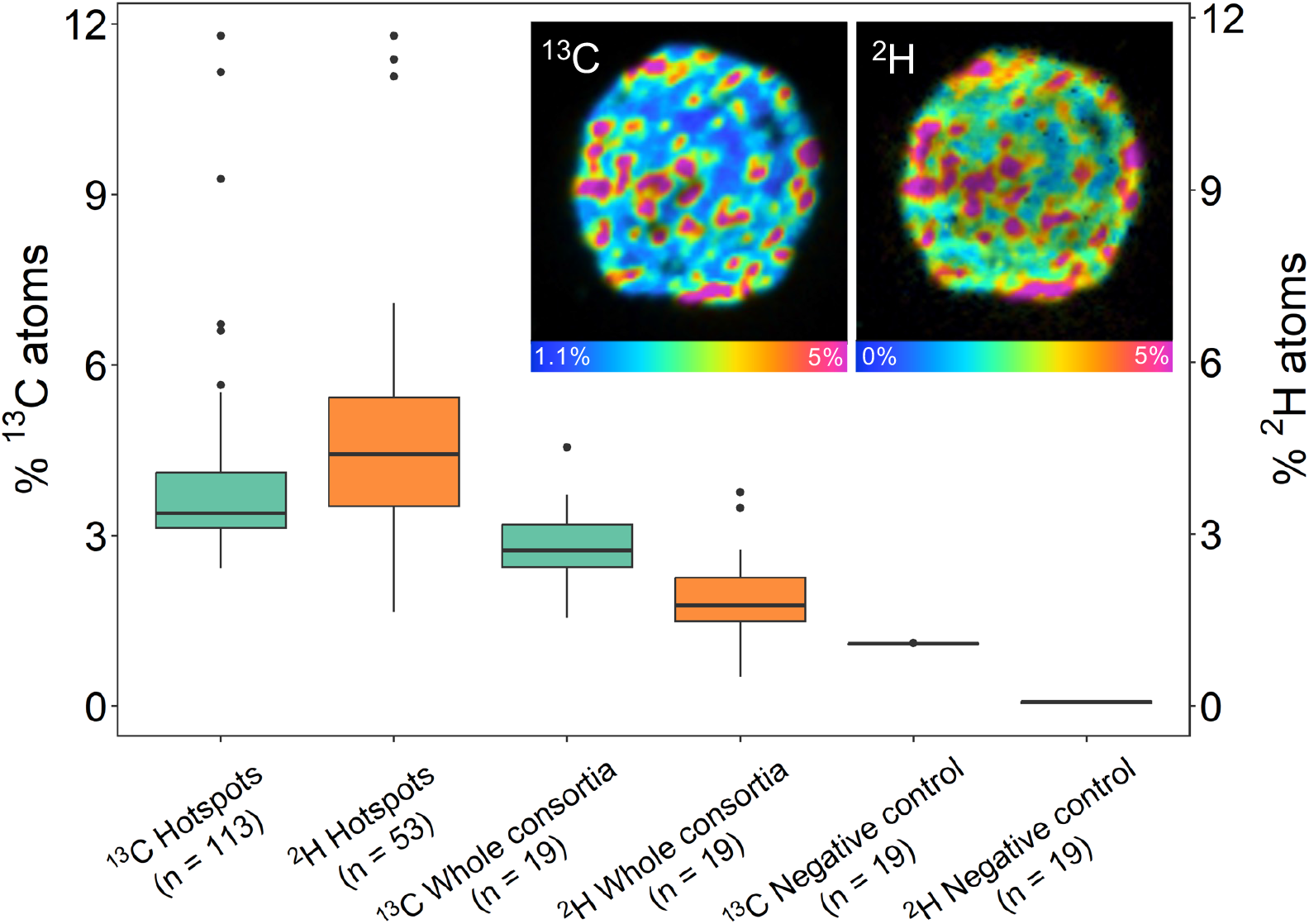
NanoSIMS analysis of MMB consortia incubated with 1,2-^13^C_2_-acetate and ^2^H_2_O. Hotspots within individual consortia were auto-segmented in ImageJ and the isotope ratios of hotspots compared to the value for the whole consortium and negative controls. The ^13^C and ^2^H hotspots showed significantly higher isotopic enrichment when compared to the values for the respective whole consortium (p <1.3x10^-3^ and <2.2x10^-8^, respectively), indicating they are metabolically differentiated. For further description of boxplots, see SI Appendix Text. Inset images show NanoSIMS HSI of the same MMB consortium analyzed using mass ratio ^13^C^12^C/^12^C_2_ and ^2^H/^1^H, revealing cell-to-cell differentiation. The HSI are scaled to show the atom percent of the respective isotope. For an example of the correlative microscopy workflow used to study MMB see Fig. S12. For ROIs, see Fig. S14.

### Metabolic differentiation as studied by BONCAT

To determine if protein synthesis was localized to specific or individual cells within the consortium, we combined BONCAT with confocal laser scanning microscopy. Our analysis revealed an apparent gradient of newly synthesized proteins within each cell of the consortium, showing localization around the acellular center of individual consortia (Fig. 7). This distinct pattern of protein synthesis was observed in all 57 MMB we examined (Fig. S16). The localization of newly synthesized protein around the acellular center of the consortium suggests this area is highly active, however the reason is currently unknown. Cells within the consortium could engage in a division of labor by metabolizing specific substrates (*e.g.*, acetate) and then sharing those resources with other cells through the acellular space, possibly by the utilization of membrane vesicles (52). A prime example of a division of labor in multicellular bacteria is the filamentous cyanobacteria Anabaena. This organism has established a mutually beneficial interaction between the heterocyst and vegetative cells via intercellular exchange of metabolites through septal junctions (5, 73). However, there is no evidence that such pores or channels exist in MMB, although an alternative route for metabolite transfer could be the acellular space within the consortium. This space has been hypothesized to be used for communication and metabolite exchange because it provides the shortest distance between any two cells (52). The localization of newly synthesized protein around the acellular center of the consortium suggests this area is highly active, possibly for exchange of metabolites from cells that are hotspots for anabolic activity. This implies cells within the consortium could metabolize specific substrates (*e.g.* acetate) and then share those resources with other cells through the acellular space, possibly by the utilization of membrane vesicles (52).

**Fig. 7.**
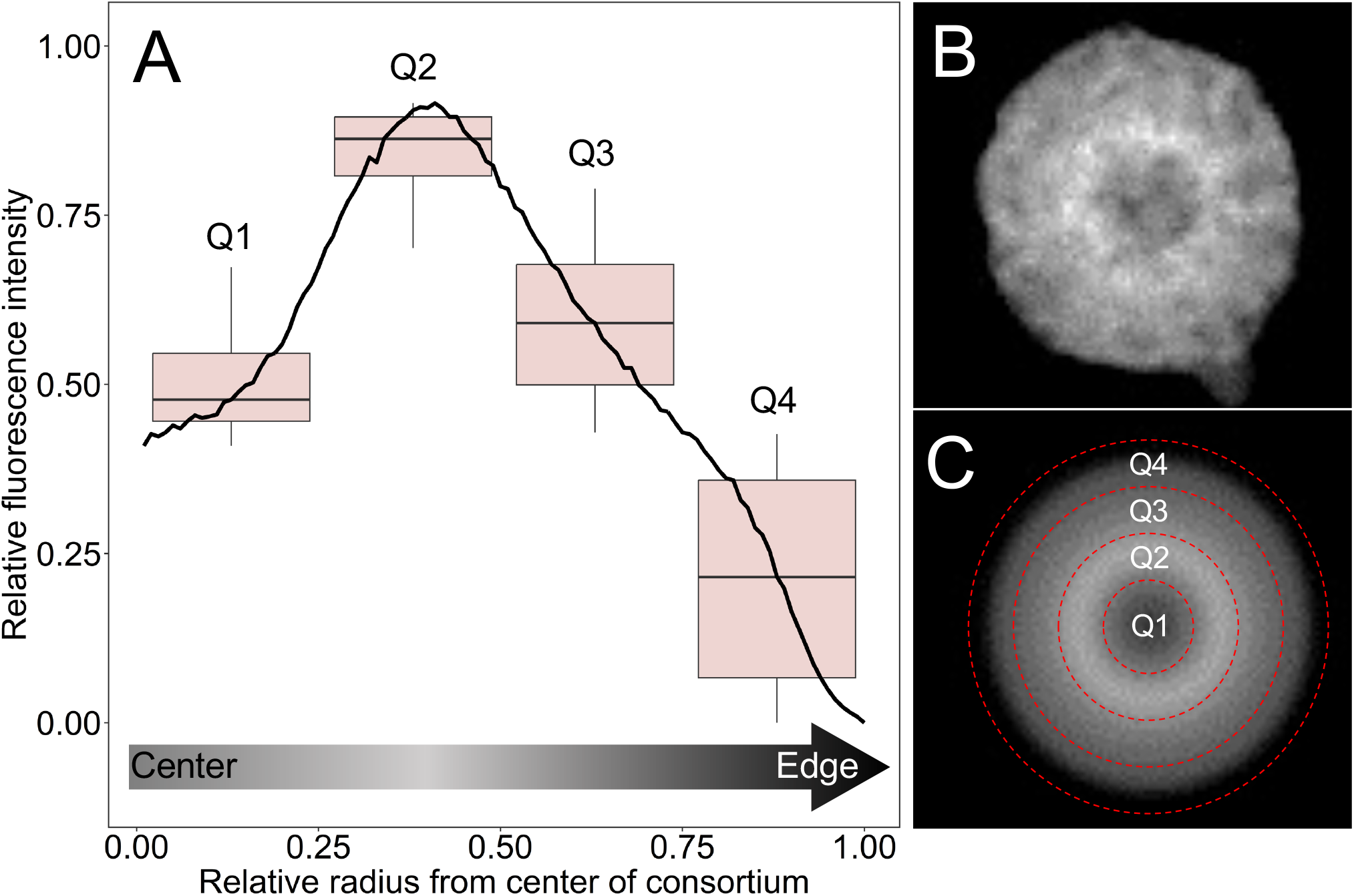
Heterogeneity in anabolic activity within individual MMB consortia as revealed by BONCAT. (*A*) The averaged intensity profile across the diameter of 57 rotationally averaged BONCAT-labeled MMB with standard deviation shown in gray. Relative fluorescence intensity (RFI) and diameter of each MMB was scaled as a ratio (0 to 1) to account for differences in fluorescence intensity between consortia and size of consortia. The boxplots show the averaged RFI for each quarter section of the radius with a pairwise statistical difference of p < 1.0x10^-10^. For further description of boxplots, see SI Appendix Text. (*B*) Gray scale confocal microscopy image of a BONCAT labeled MMB showing proteins that had been synthesized over a 24-hour period. (*C*) Image of the MMB shown in (*B*) that has been rotationally averaged prior to quantification in Eman2. The red dotted line shows each quarter analyzed for the boxplots shown in (*A*). For raw and rotationally averaged images of all 57 MMB, see Fig. S16.

## Conclusion

In summary, our study demonstrated that cutting-edge culture-independent approaches can reveal fundamental biology of yet uncultured multicellular microorganisms. We showed that MMB exhibit a higher level of complexity than previously thought by maintaining genomic heterogeneity and metabolic differentiation amongst the individual cells of a consortium. Moreover, we provided a detailed analysis of the genetic potential of eight newly discovered species of MMB as well as their ecology, ecophysiology, and *in situ* activity. We hope that these results will eventually lead to MMB representatives to be brought into culture. In addition, our results provide the basis for future experiments to further explore the mechanisms of cell-to-cell heterogeneity. Specifically, we expect mRNA-FISH (74, 75) studies to reveal to what extent gene expression levels differ from cell to cell, and SIP-NanoSIMS and spatial metabolomics (76) to reveal the molecular underpinnings of cellular interactions. Given that the biology of MMB is, as far as we know, unique in the bacterial domain, we propose MMB should, despite their recalcitrance to cultivation, receive higher attention by researchers interested in the evolution and biology of bacterial multicellularity.

## Materials and Methods

### MMB sorting, single consortia genomic sequencing and clonality analyses

A sediment sample from LSSM was shipped overnight to the Joint Genome Institute (JGI, then Walnut Creek, CA) where a magnetic enrichment was performed to obtain a pellet of MMB (see SI Appendix Methods for details). The enriched MMB were stained with SYBR Green (ThermoFisher, Eugene, OR) and sorted using a BD Influx fluorescence-activated cell sorter based on size (448 nm excitation of SYBR vs. side scatter; Fig. S17) to obtain individual MMB consortia in single wells of a 384 well plate. In addition, replicates of 10, 30, 60, and 100 cells from a culture of *Pseudomonas putida* KT2440 that had been grown in LB media were sorted into single wells as a mock control for clonal multicellularity. The *P. putida* culture liquid culture was initiated from a single colony picked from an LB agar plate. Sorted MMB and *P. putida* were then lysed and DNA amplified via the WGA-X protocol (77). Amplified SCMs were screened using 16S rRNA gene PCR according to DOE JGI standard protocols (78). Next, sequencing libraries were generated from amplified DNA using the Nextera XT v2 library preparation kit (Illumina), and sequenced on the Illumina NextSeq platform. Assemblies were derived from the IMG/M database (79). Contigs larger than 2kb were organized into genome bins based on tetranucleotide sequence composition with MetaBat2 (80) with default settings. Metagenome assembled genome (MAG) completeness and contamination were estimated with CheckM (v1.012) (81). Gene calling was performed with Prodigal (82) using the bacterial code (translation table 11). Average nucleotide identities (ANI) between MAGs were calculated with FastANI (v1.1) (83), filtered at 95% sequence identity and 30% aligned fraction, and then clustered using mcl (v14-137) (84).

We assessed clonality of sorted MMBs, single sorted and amplified *Pseudomonas* controls and other MAGs derived from sorted MMBs by mapping the reads from the respective libraries to the contigs larger than 5kb in assemblies derived from the same library using BBMap (v38.79) (https://sourceforge.net/projects/bbmap/, (85)) with the flags minid=0.95 minaveragequality=30. Variants were called with the BBTools script callvariants.sh using the flags minreads=2 minquality=30 minscore=30 minavgmapq=20 minallelefraction=0.05 and identified variants were then annotated as synonymous (s), nonsynonymous (ns) or intergenic depending on their position. Variants made up by one or more Ns were excluded from the analysis. To investigate differences between MMB, all libraries were also mapped to contigs with a size of at least 5kb infrom the longest MMB assembly (3300034493).

### Stable isotope probing

To empirically test the use of carbon substrates as predicted by the functional annotation of MMB SCMs and determine the anabolic activity of MMB cells, we employed performed both in situ and in vitro incubations of MMB with ^13^C- and ^2^H-labeled substrates (all 99.9%, Cambridge Isotopes Laboratories). The *in situ* incubations were performed in duplicate on August 28^th^ 2022 at LSSM by amending 200 mL top sediment slurry with 2 mM ^13^C-1,2-acetate, 2 mM ^13^C-1,2- succinate, 5 mM ^13^C-1,2-propionate, 5 mM ^13^C-bicarbonate, or 2 mM ^13^C-1,2-acetate plus 8 mM molybdate (a competitive inhibition of sulfate reduction). A negative control to which no amendment was made as well as a killed control in which biomass had been pre-incubated with 4% paraformaldehyde (PFA) for 60 minutes at ambient temperature prior to addition of 2 mM ^13^C- 1,2-acetate were also performed. Samples were stored in 200 mL Pyrex glass bottles (Corning, Glendale, AZ) and incubated for 24 hours *in situ* at the sample site where they were buried 4-6 cm below the sediment in a basket (Fig. S1C-D). The *in vitro* incubations were performed by incubating magnetically enriched MMB in 10 mL of 0.22 µm filter sterilized (Millipore, Burlington, MA) LSSM water amended with the same amendments as the *in situ* incubations, as well as 50% deuterium oxide (D2O), for 24 hours at ambient lab temperature (∼23 °C) in the dark. At the end of each incubation period, MMB were magnetically enriched and fixed with 4% PFA for 60 minutes at ambient temperature. Cells were centrifuged for 5 minutes at 16,000 g, after which the supernatant was removed, and the cell pellets resuspended in 50 µL 1× PBS and stored at 4 °C.

### NanoSIMS

Samples were prepared for NanoSIMS on stainless steel coupons as previously described (86); for details see SI Materials and Methods. To quantify cell-to-cell differences in isotope uptake within individual consortia, ROIs were selected around localized densities (*i.e*., hotspots) of masses corresponding to the respective substrate and compared to whole consortia values for the same isotope of interest. To select ROIs, Fiji (https://imagej.net/software/fiji/) was used to convert the mass image to an 8-bit image for which the brightness and contrast adjusted to help identify the localized densities for the mass of interest (*e.g*. ^12^C^2^H 14.02, ^12^C^13^C 25.00).

### BONCAT

BONCAT was performed as previously described (87); for details see SI Materials and Methods. To evaluate cell-cell differences in anabolic activity of individual consortia, MMB were imaged by taking z-stacks (approximately 300 nm per image) of the entire consortia using an Inverted DMI8 Stellaris 8 Confocal Microscope (Leica Microsystems). Images focused on the center of the consortia were selected and Eman2 (88) was used to select individual MMB for particle analysis. Each image was then filtered using an edge mean normalization, center of mass xform, and rotational average math settings (Fig. S16). Because of varying sizes of consortia, a Python script was used to determine the radius of each consortium by calculating the number of pixels from the center of mass, as determined by the filter, to where the standard deviation of the pixels is < 0.01. The radius of all consortia was standardized by dividing 1 by the radius. Additionally, the average fluorescence intensity was normalized by calculating *I*_*norm*_ = 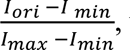, where *I*_*ori*_ is the original fluorescence intensity value and *I*_*min*_/*I*_*max*_ are the minimum and maximum relative fluorescence intensity values for the individual consortia. The average and standard deviation of data was calculated and plotted using R. All code used for analysis is deposited on GitHub (https://github.com/georgeschaible/MMB-BONCAT).

## Supplementary Methodology

Sample collection, phylogenetic analysis, genome and magnetosome analyses, FISH, BONCAT, NanoSIMS, Raman microspectroscopy, and SEM experiments, geochemical analysis, and statistical analyses are described in the SI Materials and Methods.

## Acknowledgements

This study was funded through NASA Exobiology program award NNX17AK85G to RH and NASA FINESST award 80NSSC20K1365 to GS and RH. CG was supported by the National Institute of General Medical Sciences (P30GM140963). SER was supported by the Simons Foundation (824763). A portion of this research was performed under the Community Sciences Program (awards DOI: 10.46936/10.25585/60001107 and DOI: 10.46936/10.25585/60001212) and used resources at the DOE Joint Genome Institute (https://ror.org/04xm1d337), which is a DOE Office of Science User Facility operated under Contract No. DE-AC02-05CH11231. A portion of this research was performed under the Facilities Integrating Collaborations for User Science (FICUS) program (awards DOI: 10.46936/fics.proj.2017.49972/6000002 and 10.46936/fics.proj.2020.51544/60000211) and used resources at the Environmental Molecular Sciences Laboratory (https://ror.org/04rc0xn13), which is a DOE Office of Science User Facilities operated under Contract No. DE-AC05-76RL01830. This work was performed in part at the Montana Nanotechnology Facility, an NNCI member supported by NSF grant ECCS-2025391. Fluorescence and Raman microscopy imaging was made possible by The Center for Biofilm Engineering Imaging Facility at Montana State University, which is supported by funding from the NSF MRI Program (2018562), the M. J. Murdock Charitable Trust (202016116), the US Department of Defense (77369LSRIP), and by the Montana Nanotechnology Facility (an NNCI member supported by NSF Grant ECCS-2025391). Montana State University’s Confocal Raman microscope was acquired with support by the National Science Foundation (DBI-1726561) and the M. J. Murdock Charitable Trust (SR-2017331). We thank Jeffrey Marlow (Boston University), Rachel Spietz (MSU), and Ashley Cohen (MSU) for help with the collection of LSSM sediment samples and assistance with lab work as well as Heidi Smith (MSU) for microscopy support. We also thank Anthony Kohtz, Amanda Wilkins, and Hope McWilliams (all MSU) for assistance with lab work, Marike Palmer (University of Nevada Las Vegas) for discussions on taxonomy, Julie Huber (Woods Hole Oceanographic Institution) for graciously providing access to her lab space at WHOI, and Kristina Hillesland (University of Washington, Bothell) for critical comments that helped to improve the manuscript. We thank our Brazilian colleagues Fernanda Abreu, Henrique Lins de Barros, Marcos Farina, Carolina Keim, and Juliana Martins (Lopez), as well as Sherri Simmons for their foundational work on MMB and allowing us to name newly discovered MMB species after them and in honor of the late Ulysses Lins, who transformed our understanding of MMB.

## Data availability

The single consortia metagenomes of MMB generated in this study are available on JGI’s IMG/M under the genome numbers 3300028595, 3300034483-3300034486, and 3300034488- 3300034505. The genome sequences of *Ca*. M. multicellularis and *Ca*. M. HK-1 are available at NCBI Genbank under accession numbers GCA_000516475 and JPDT00000000, respectively. Magnetosome sequences for *Ca*. Desulfamplus magnetomortis BW-1, *Ca*. Magnetananas rongchenensis RPA, and MMP XL-1 are available at GenBank under accession numbers HF547348, KY084568, and ON204283:ON204284, respectively. Python and R code used to analyze BONCAT data are available on GitHub (https://github.com/georgeschaible/MMB-BONCAT).

## Supporting Information

### SI Results and discussion

#### Protologue

We assign type genomes for eight newly discovered species of MMB and propose the following provisional taxonomic assignments. All researchers were contacted and gave permission to name new MMB species after them. See SI Appendix Table S4.

- *Candidatus* Magnetoglobus abreuianus sp. nov. a.bre.u.i.a’nus N.L. masc. adj. abreuianus; named in honor of Fernanda Abreu, who described the first species of MMB, Magnetoglobus multicellularis (1). This uncultured species is represented by bin 3300034485, which has an estimated completeness of 89.22%, a contamination of 2.09%, with no 16S rRNA, 23S rRNA or 5S rRNA genes.
- *Candidatus* Magnetoglobus debarrosii sp. nov. de.bar.ro’si.i N.L. gen. n. debarrosii, of de Barros; named in honor of Henrique Lins de Barros, who shaped understanding of MMB for the past four decades. This uncultured species is represented by bin 3300034500, which has an estimated completeness of 90.62%, a contamination of 1.94%, and contains 16S rRNA, 23S rRNA and 5S rRNA genes.
- *Candidatus* Magnetoglobus farinai sp. nov. fa.ri.’na.i N.L. gen. n. farinai, of Farina; named in honor of Marcos Farina, who co-discovered MMB in 1983 (2, 3). This uncultured species is represented by bin 3300034494, which has an estimated completeness of 90.65%, a contamination of 0.86%, and contains 16S rRNA, 23S rRNA and 5S rRNA genes.
- *Candidatus* Magnetoglobus keimiae sp. nov. ke.i’mi.ae N.L. gen. n. keimiae, of Keim; named in honor of Carolina Keim, who first demonstrated the multicellular life cycle of MMB (4). This uncultured species is represented by bin 3300034495, which has an estimated completeness of 94.77%, a contamination of 1.53%, and contains 16S rRNA, 23S rRNA and 5S rRNA genes.
- *Candidatus* Magnetoglobus linsii sp. nov. lin’si.i N.L. gen. n. linsii, of Lins; named in honor of the late Ulysses Lins, whose pursuit of pure, “romantic” scientific questions (5) shaped our understanding of MMB. This uncultured species is represented by bin 3300034496, which has an estimated completeness of 93.56%, a contamination of 1.29%, and contains 16S rRNA, 23S rRNA and 5S rRNA genes.
- *Candidatus* Magnetoglobus martinsiae sp. nov. mar.tin’si.ae N.L. gen. n. martinsiae, of Martins; named in honor of Juliana Lopes Martins’ contributions to the study of MMB. This uncultured species is represented by bin 330034493, which has an estimated completeness of 91.77%, a contamination of 0.86%, with no 16S rRNA, 23S rRNA or 5S rRNA genes.
- *Candidatus* Magnetoglobus simmonsiae sp. nov. sim.mon’si.ae N.L. gen. n. simmonsiae, of Simmons; named in honor of Sherri Simmons, whose research on MMB in Little Sippewissett Salt Marsh laid the foundation for much of our analysis (6). This uncultured species is represented by bin 3300034505, which has an estimated completeness of 85.37%, a contamination of 1.31%, and contains 16S rRNA, 23S rRNA and 5S rRNA genes.
- *Candidatus* Magnetomorum sippewissettense sp. nov. sip.pe.wis.set.ten’se N.L. neut. adj. sippewissettense; pertaining to Sippewissett, named after Little Sippewissett Salt Marsh, Falmouth, MA, USA, where this study was conducted. This uncultured species is represented by bin 3300034504, which has an estimated completeness of 86.63%, a contamination of 0.32%, and contains 16S rRNA, 23S rRNA and 5S rRNA genes.

### Failure to establish an enrichment culture

Previous studies have attempted to cultivate magnetically enriched MMB in defined media but so far there has been no success despite the ability to magnetically enrich them to >99% purity (7, 8). In an attempt to bring MMB into cultivation, we designed a medium (SI Appendix Table S12) informed by the geochemical composition of the water at LSSM (SI Appendix Table S13), the metabolic predictions derived from genomic data (Fig. 4; SI Appendix Table S7), and the results of SIP-NanoSIMS experiments (Fig. 5; SI Appendix Table S10). Incubations were performed under anoxic conditions at 27 °C and a pH of 7.4. MMB were found to maintain their magnetotaxis and could be recovered from the media for up to 15 days, after which no MMB could be magnetically enriched nor identified using FISH.

### Characterization of magnetosome and light sensing genes

Previous spectroscopic analysis has indicated the utilization of greigite magnetosomes in LSSM MMB (9), though genes relating to greigite production in LSSM MMB have not previously been identified. Genomic analysis of MMB from other locations (i.e., German Wadden Sea) revealed they are capable of synthesizing magnetite and/or greigite within their magnetosome, though greigite is most common due to environmental thermodynamic restrictions (10–13). We identified core greigite biomineralization genes in all single consortia metagenomes (SCMs) (*mamA*, B*, E-Cter*, E-Nter*, I-4*, I-5*, MB-like*, O*, Q*,* and *T\** as well as *mad12, 14, 17-19, 23-30,* and *mamK)* and magnetite biomineralization genes in SCM 3300034500. The organization of the magnetosome gene clusters (MGCs) was conserved across LSSM SCMs. The synteny of the greigite biomineralizing genes were similar to *Ca*. Magnetoglobus multicellularis and MMP XL- 1, although *Ca*. M. sippewissettense appears to lack the organization found in *Ca*. *Magnetoglobus* species. The synteny of magnetite biomineralizing genes in 3300034500 was conserved across *Ca*. Magnetomorum HK-1, *Ca*. Magnetananas rongchenensis RPA, MMP XL-1, and *Desulfamplus magnetomortis* BW-1 (Fig. S18). Greigite and magnetite synthesizing genes have been identified in the genomes of aforementioned MMB but greigite appears to be preferentially used over magnetite (10, 11), which is congruent with observations of LSSM MMB (Fig. S7). An explanation for the presence of magnetite biomineralizing genes in 3300034500 could be horizontal gene transfer (14), although their function/role in the environment is unclear. The SCM MGCs contained additional genes surrounding the core greigite magnetosome genes including genes encoding for actin-related proteins, rod shape-determining protein MreB, and chemotaxis protein CheF, all potentially involved in the formation and maintenance of the magnetosome (Fig. S18; SI Appendix Table S14).

Genomic and *in vitro* observations indicate light plays an important role in the behavior and position of MMB in the sediment column and has even been shown to be responsible for triggering cell division (7, 15, 16). The *kaiB* and *kaiC* genes, involved in circadian cycle, and genes for bacteriophytochrome and photoactive yellow protein were recovered from the SCMs (SI Appendix Table S7), supporting previous observations of LSSM MMB response to light (15). In addition, multiple copies of two-component chemotaxis genes were identified in the SCMs. The combination of genes related to magnetotaxis, phototaxis, and chemotaxis likely enables MMB to effectively navigate environmental gradients. Moreover, the identification of genes protecting against oxygen radicals (SI Appendix Table S7) implies MMB are potentially capable of survival in (micro)oxic sediment layers. Taken together, our finding suggests LSSM MMB likely maintain constant movement along chemical gradients in their surroundings, as has been previously suggested (7).

## SI Materials and methods

### Sample collection and magnetic enrichment of MMB

Sediment samples were collected from a tidal pool at Little Sippewissett salt marsh (LSSM, 41.5758762, -70.6393191) in Falmouth, MA (USA) during low tide on October 2^nd^ 2018, August 17^th^ 2020, September 21^st^ 2021, and August 28^th^ 2022. For each sample, 1 L of sediment slurry (7:3 sediment to water ratio) was collected in plastic bottles and shipped within one day on ice to Montana State University, Bozeman, MT (USA), where the slurry was transferred to a 1 L glass beaker and stored in the dark at ambient laboratory temperature (∼23°C). MMB were magnetically enriched from the sediment by placing the South end of a magnetic stir bar against the exterior of the glass beaker just above the sediment layer, agitating the sediment by stirring, and then allowing the sediment to settle for 60 minutes. Magnetically enriched MMB were collected by pipette and further enriched as previously described (9) (SI Video 1).

### Scanning electron microscopy (SEM) and cellulase experiment

To acquire SEM micrographs of MMB, a Zeiss (Jena, Germany) SUPRA 55VP field emission scanning electron microscope (FE-SEM) was operated at 1 keV under a 0.2–0.3 mPa vacuum with a working distance of 5 mm and 30 μm aperture. For the cellulase experiment, samples of magnetically enriched MMB were incubated for 1 hr at 37°C in 0.22 µm filtered LSSM water with a pH adjusted to 5 for optimal cellulase activity. MMB were treated with 5 mg/mL of cellulase (MP Biomedicals, Solon, OH USA) as per the manufacturer’s instructions. A control reaction under the same conditions but without cellulase was performed to check the effect of temperature and low pH on MMB. The incubation was stopped by the addition of PFA to a final concentration of 4% and samples incubated at ambient temperature for 1 hr, after which cells were centrifuged at 16,000 g for 5 minutes and the supernatant removed, and cells resuspended in 1x PBS. Cells were dried onto a mirrored stainless-steel slide and dried at 46 °C for 2 minutes, after which they were washed in MilliQ water three times for 10 seconds each and the slide was air dried. All electron microscopy work was performed at the Imagining and Chemical Analysis Laboratory (ICAL) of Montana State University (Bozeman, MT). No conductivity coating was applied prior to analysis.

### Phylogenetic, Phylogenomic, and Comparative Genomic analyses

The 16S rRNA gene sequences encoded in the MMB SCMs were used in BLASTn (17) searches to screen the NCBI database for related sequences (SI Appendix Table S2). All 16S rRNA sequences were aligned using SSU-ALIGN and a maximum likelihood analysis was performed using FastTree2.1 with 500 ultrafast bootstraps (18, 19). Of 139 single-copy bacterial genes searched (20), a subset of six were present in all 22 SCM (SI Appendix Table S3). These were aligned with reference sequences using MUSCLE (21), concatenated, and phylogenetically analyzed with FastTree2.1 (500 ultrafast bootstraps) (18). Average nucleotide identities (ANIs) of SCMs and 16S rRNA sequences were calculated with FastANI (22) and pairwise BLASTn comparisons, respectively.

### Genome annotation

The metabolic potential of MMB SCMs was determined by mapping gene annotations provided by IMG/M (23) to metabolic pathways outlined in the KEGG (Kyoto Encyclopedia of Genes and Genomes) database (24). Further investigation of genes was done by inspection of gene neighborhoods and identification of conserved domains and motifs through submission of genes to the NCBI conserved domain database (25) and MPI Bioinformatics HHpred Toolkit (26). Classification of hydrogenases was done using HydDB (27) and if a subunit is membrane bound or soluble determined using DeepTMHMM (28).

### Comparative genomic analysis of magnetosome gene clusters

To identify the magnetosome gene clusters, pairwise BLASTn comparisons of individual magnetosome genes from *Ca*. Desulfamplus magnetomortis BW-1 (HF547348) (29) were performed on each of the individual SCMs as well as the reference genomes of *Ca*. Magnetoglobus multicellularis (IMG ID 2558860350) (7) and *Ca*. Magnetomorum sp. HK-1 (IMG ID 2648501189) (11). Gene synteny figures of magnetosome encoding loci were made with Clinker (v0.0.27) using default settings and an identity setting of 0.45 (30).

### Fluorescence *in situ* hybridization (FISH)

Double-labeled oligonucleotide probes for FISH (DOPE-FISH, (31)) were purchased from Integrated DNA Technologies (Coralville, IA) to visualize different MMB taxa. Genus level populations of MMB were targeted by using newly designed DOPE-FISH probes targeting the 1032-1049 nt region of the 16S rRNA (*E. coli* equivalent) using full length 16S rRNA gene sequences from the MMB SCMs and previously published 16S RNA gene clone sequences from LSSM (6) and the two reference genomes. Probes were designed to target five genus level populations of MMB in LSSM (groups 1-5) as well as three individual species within groups 1 and 2 (Fig. S9; SI Appendix Table S9). Probes were designed manually using ARB (32) and evaluated *in silico* using the TestProbe tool of Silva ((33), http://arb-silva.de, database release 138.1), the MatchProbe tool of ARB, and mathFISH ((34), http://mathfish.cee.wisc.edu/). All probes have at least one central mismatch to non-target sequences (SI Appendix Table S9) and were verified in the Silva database (33). To ensure stringency of each probe, competitor probes were designed for each probe and used accordingly. Group-specific probes were designed to compete for the same binding site to guarantee specific binding. Specificity of genus-specific probes was checked using hybridization curve assays in CloneFISH (35) experiments using representative sequences for each of the five MMB groups. Fixed cells were dehydrated using an increasing ethanol series (1 min in each 50, 80, and 96% ethanol) and FISH was carried out on Teflon coated glass slides. Samples were hybridized for three hours in a humid chamber at 46 °C with a final probe concentration of 2.5 ng μL^−1^. Positive and negative controls using EUB338 and NonEUB338 (36) were conducted routinely. Neither in CloneFISH nor in environmental FISH experiments, *E. coli* cells or MMB, respectively, were labeled by more than one MMB group- or species-specific probe, demonstrating specificity of the newly designed probes at the final formamide concentrations (SI Appendix Table S9).

### Bioorthogonal noncanonical amino acid tagging (BONCAT) and confocal fluorescence microscopy

To evaluate the activity of MMB within LSSM, BONCAT incubations were performed on LSSM sediments. A 15 cm long sediment core was collected on August 17^th^ 2020 from the West end of the sample site and shipped to MSU overnight. Upon receipt, the core was sectioned into 1 cm horizons that were homogenized and divided into triplicate 25 mL serum vials. Vials were placed in an anoxic chamber (Coy Lab Products, Grass Lake, MI) and 10 mL of 0.22 µm filtered LSSM water (made anoxic by bubbling with nitrogen gas for 60 minutes) added to each vial. Samples were amended with 50 μM L-Homopropargylglycine (HPG, Click Chemistry Tools, Scottsdale, AZ) except for triplicate negative controls. Samples were incubated for 24 hours in the dark at ambient lab temperature, after which MMB were magnetically enriched from each triplicate horizon incubation and fixed in 4% PFA. Cells were centrifuged for 5 minutes at 16,000 g, after which the supernatant was removed, and the cell pellets resuspended in 50 µL 1× PBS and stored at 4 °C. To fluorescently label alkyne-tagged proteins, cells were dried to a glass slide and dehydrated using an ethanol series (50, 80, and 96% for three minutes each). Click chemistry using AlexaFlour-405-Azide was performed according to published methods (37). In addition, DOPE-FISH was performed on the samples to identify individual Groups of MMB (see SI). Cells were imaged using a Leica DM4B epifluorescent microscope (Leica Microsystems, Deerfield, IL USA) and relative fluorescence intensity calculated using Daime with normal edge thresholding settings (38).

To evaluate differences in activity within individual MMB consortia, sediments containing MMB were amended with 50 µM *L*-azidohomoalanine (AHA, Click Chemistry Tools, Scottsdale, AZ USA) and incubated at ambient temperature in the dark for 24 hours, after which the MMB were magnetically enriched and fixed in 4% PFA for 60 minutes at ambient temperature. Cells were centrifuged for 5 minutes at 16,000 g, after which the supernatant was removed, and the cell pellets resuspended in 50 µL 1× PBS and stored at 4 °C. To fluorescently tag azide-labeled proteins, cells were dried to a glass slide and dehydrated using an ethanol series (50, 80, and 96% for three minutes each). Click chemistry using AlexaFlour-488-Alkyne was performed using published methods (37).

### Confocal Raman microspectroscopy and spectral processing

Raman spectra of individual MMB were acquired using a LabRAM HR Evolution Confocal Raman microscope (Horiba Jobin-Yvon) equipped with a 532 nm laser and 300 grooves/mm diffraction grating. Spectra of the MMB were acquired using a 100× dry objective (NA = 0.9), with 10 acquisitions of 2 seconds each, and a laser power of 4.5 mW. Spectra were processed using LabSpec version 6.5.1.24 (Horiba) with a Savitsky-Goly smoothing algorithm, baselined, and finally normalized to the maximum intensity within the 2,800-3,100 cm^-1^ regions. Peaks corresponding to lipids, PHB, and exopolysaccharides were identified in previous studies (39, 40) and are listed in SI Appendix Table S8.

### NanoSIMS

Ion images were acquired using the NanoSIMS 50L (Cameca) at the Environmental Molecular Sciences Laboratory at the Pacific Northwest National Laboratory. All NanoSIMS images were acquired using a 16 keV Cs+ primary ion beam at 512 × 512-pixel resolution with a dwell time of 13.5 ms px^−1^. Analysis areas were pre-sputtered with ∼ 1016 ions cm^−2^ prior to analysis. Secondary ions were accelerated to 8 keV and counted simultaneously using electron multipliers (EMs). The vacuum gauge pressure in the analytical chamber during all analyses was consistently less than 3 × 10^−10^ mbar. Other analytical conditions included a 200 μm D1 aperture, 30 μm entrance slit, 350 μm aperture slit, and 100 μm exit slits. The OpenMIMS plugin for ImageJ was used to access and correct images pixel by pixel for dead time (44 ns) and QSA (β = 0.5). HSI images shown in main text are filtered with a median filter ratio radius of 0.5. This filter is used to improve contrast but does not adversely affect quantitative data reported in tabular form for the regions of interest (ROIs). Data from regions of interest (ROIs) were exported to a custom spreadsheet for data reduction. Quantitative ^13^C^12^C/^12^C2 analyses were calibrated against an in-house yeast reference material of known natural abundance δ^13^C during the same analytical session using similar conditions to those used to analyze the bacterial culture samples. An unknown background signal interfering with the ^2^HC signal was subtracted using the yeast ion images but no attempt was made to calibrate the ^2^HC/^1^HC. These data are therefore not strictly quantitative, but this does not change interpretation of the relatively higher ^2^H content of the enriched samples compared with controls (Schaible, Cliff, *et al.*, manuscript in preparation). The yeast reference material had been stored in the NanoSIMS under high vacuum for several months prior to the analyses reported here. During ^2^HC/^1^HC analyses, detectors collecting secondary ^2^HC and ^1^HC ions were situated near the center of the magnet radius and Helmholtz steering coils were carefully adjusted to improve simultaneous secondary centering characteristics. Propagation of uncertainty includes counting statistics and external precision of isotopic ratios of 16 individual yeast cells.

### Geochemical analysis

Overlaying water from LSSM was collected and 0.22 μm filtered into 50 mL tubes for ion chromatography and inductively coupled plasma optical emission spectroscopy (ICP-OES). Trace-metal grade HNO3 was added to the ICP-OES tubes for a final concentration of 2%. Samples for total organic carbon (TOC) were collected by 0.22 μm filtering LSSM water into ashed glass vials. All geochemical measurements were made in the Environmental Analytical Laboratory at Montana State University (Bozeman, Montana). Details on how chemical analyses were performed can be found in Lynes, Krukenberg et al 2023 (41).

### Statistical analysis

All datasets were analyzed in R (42) using the tidyverse, rstatix, and ggpubr packages (43, 44). Statistical differences between multiple variables were determined using ANOVA and pairwise t-tests with a Bonferroni p-adjusted method. Boxplots show the distribution of the dataset, where the box corresponds to the interquartile range (IQR) containing the middle 50% of the data, the black line inside the box represents the median, and the whiskers extend to the minimum and maximum values within 1.5 times the IQR from the first and third quartiles, respectively.

### Detailed author contributions

GS and RH developed the research project and designed experiments, with input from JC on NanoSIMS analyses. GS, ER, VE, and RH collected field samples. GS conducted all wet lab experiments except NanoSIMS measurements, which were performed by JC. ZJJ and FS processed metagenomic data, assembled SCMs, and performed similarity comparisons as well as SNP and clonality tests. GS performed genome annotations. GS and ZJJ constructed phylogenies and ANIs. DG performed FACS and whole genome amplification experiments supervised by RRM. GS and CG processed and analyzed BONCAT image data. GS performed all statistical analyses and made the figures. GS and RH were responsible for funding and designed FISH probes. RH supervised the project. GS and RH wrote the manuscript draft, which was then edited by all authors.

## Video legends

**Video S1.** Third and final round of a magnetic enrichment of MMB. MMB swim from the bottom of the microcentrifuge tube up towards the magnetic North of a magnetic stir bar. Video is 30x the original speed.

**Video S2.** MMB swimming at the edge of a hanging water droplet towards the magnetic North of a magnetic stir bar that is out of frame. The temporal response of MMB to changes in the magnetic field is observed when the magnet is turned. This switches the magnetic field and induces a change in the swimming direction of the MMB consortia, until the magnet is turned again, and the MMB swim back to the edge of the hanging water droplet. Video is at its original speed.

---

**Fig. S1.**
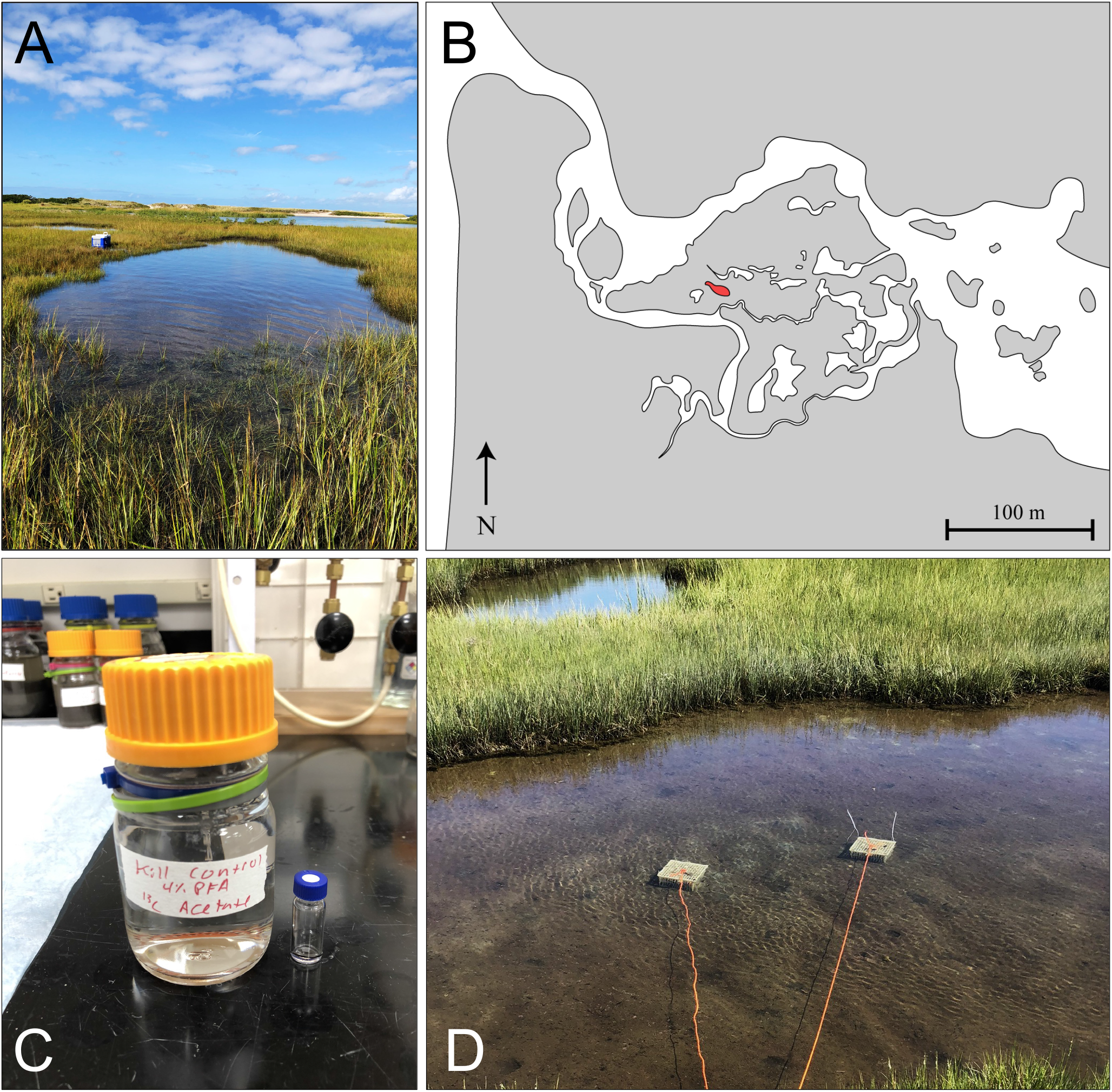
Little Sippewissett salt marsh, Falmouth MA. (*A*) Photo of the tidal pool from which sulfidic sediments were obtained, facing west towards Buzzards Bay. (*B*) Map of the salt marsh showing the tidal pool in red and water in white. (*C*) Each sample was incubated in a 200 mL bottle filled to the top with the sediment slurry and tightly capped. Because no MMB could be recovered post-fixation from the the kill control sample, 200 μL of sample were incubated in a small glass vial inside of the 200 mL bottle. (*D*) Samples were incubated *in situ* below the sediment at the site for 24 hours.

**Fig. S2.**
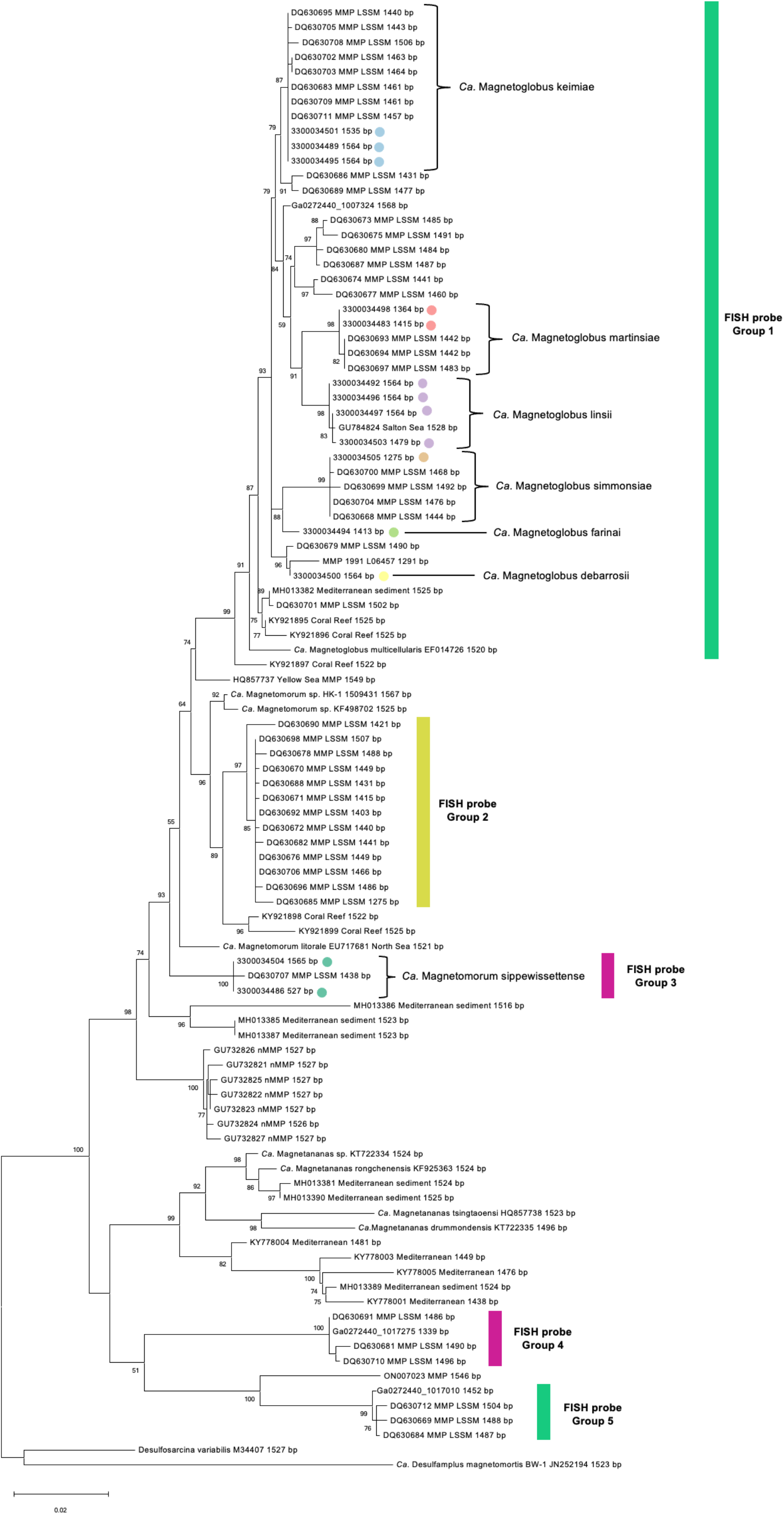
Phylogenetic analysis of MMB using near-full length 16S rRNA genes (length listed next to name) found in 14 of the 22 SCMs and in reference genomes. Tree reconstructed using maximum likelihood method with bootstrap values calculated using 500 replicates. Bootstrap values above 50 are shown. *Ca.* M. abreuianus is not shown in this analysis because no 16S rRNA gene was recovered from the SCM. Color coded sequences belong to their respective SCM, as shown in supplemental table 1. Bars on right show specificities of our newly designed FISH probes that target genus-level groups of MMB m LSSM (SI Appendix Table S9).

**Fig. S3.**
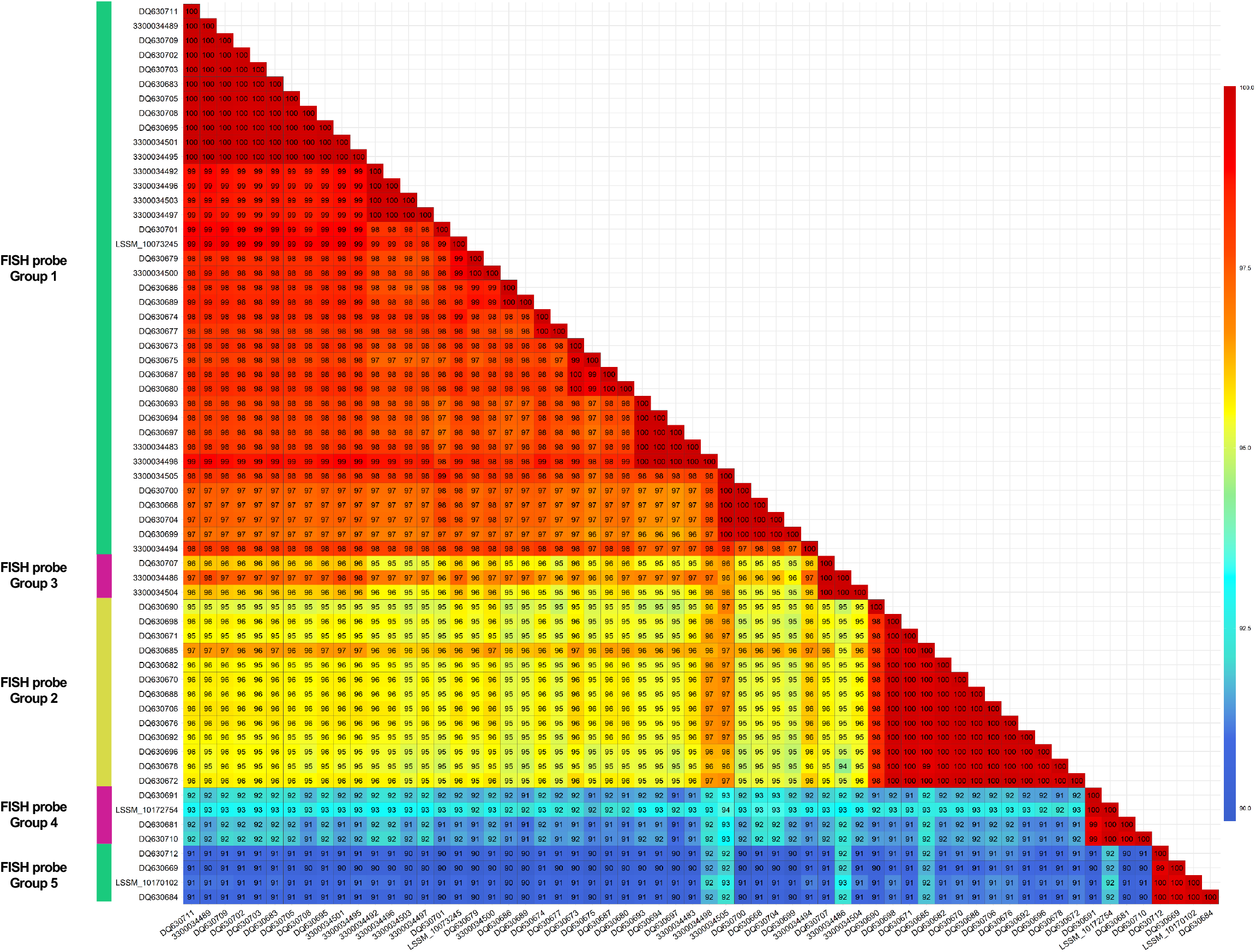
Near-full length 16S rRNA gene comparison of all sequences recovered in this study and previous studies at LSSM (Simmons and Edwards 2007). Percent identity values are shown within boxes. Bars on left highlight MMB groups for which genus-level FISH probes designed (SI Appendix Table S9).

**Fig. S4.**
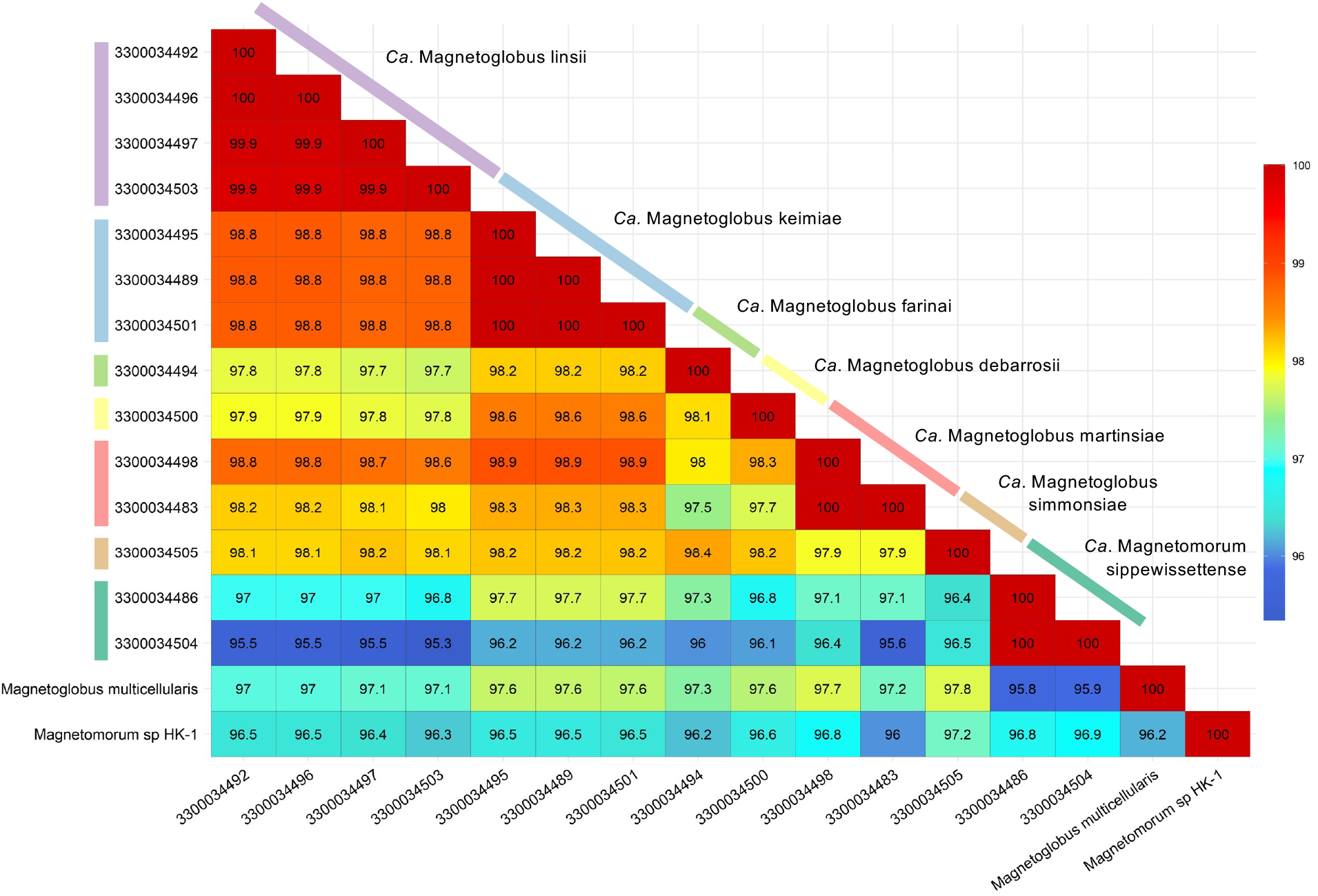
Near-full length 16S rRNA identity comparison for the 14 sequences recovered from SCMs and the two MMB reference genomes (*Ca.* M. multicellularis and *Ca.* Magnetomorum sp. HK-1). Percent identity values are shown within boxes.

**Fig. S5.**
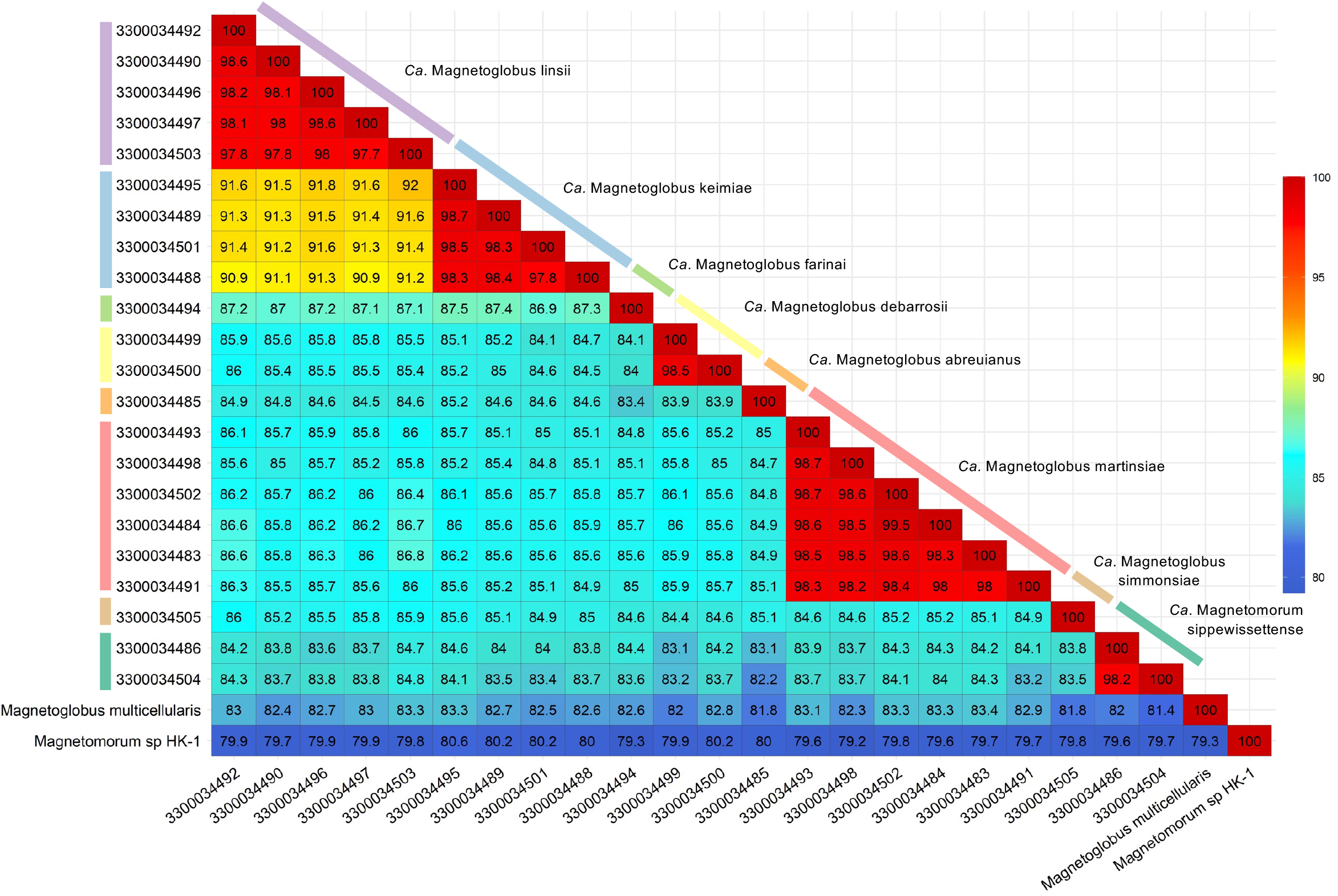
Genome ANI comparing each of the 22 individual SCMs with the two publicly available reference genomes (*Ca.* M. multicellularis and *Ca.* Magnetomorum sp. HK-1). ANI values are shown within boxes.

**Fig. S6.**
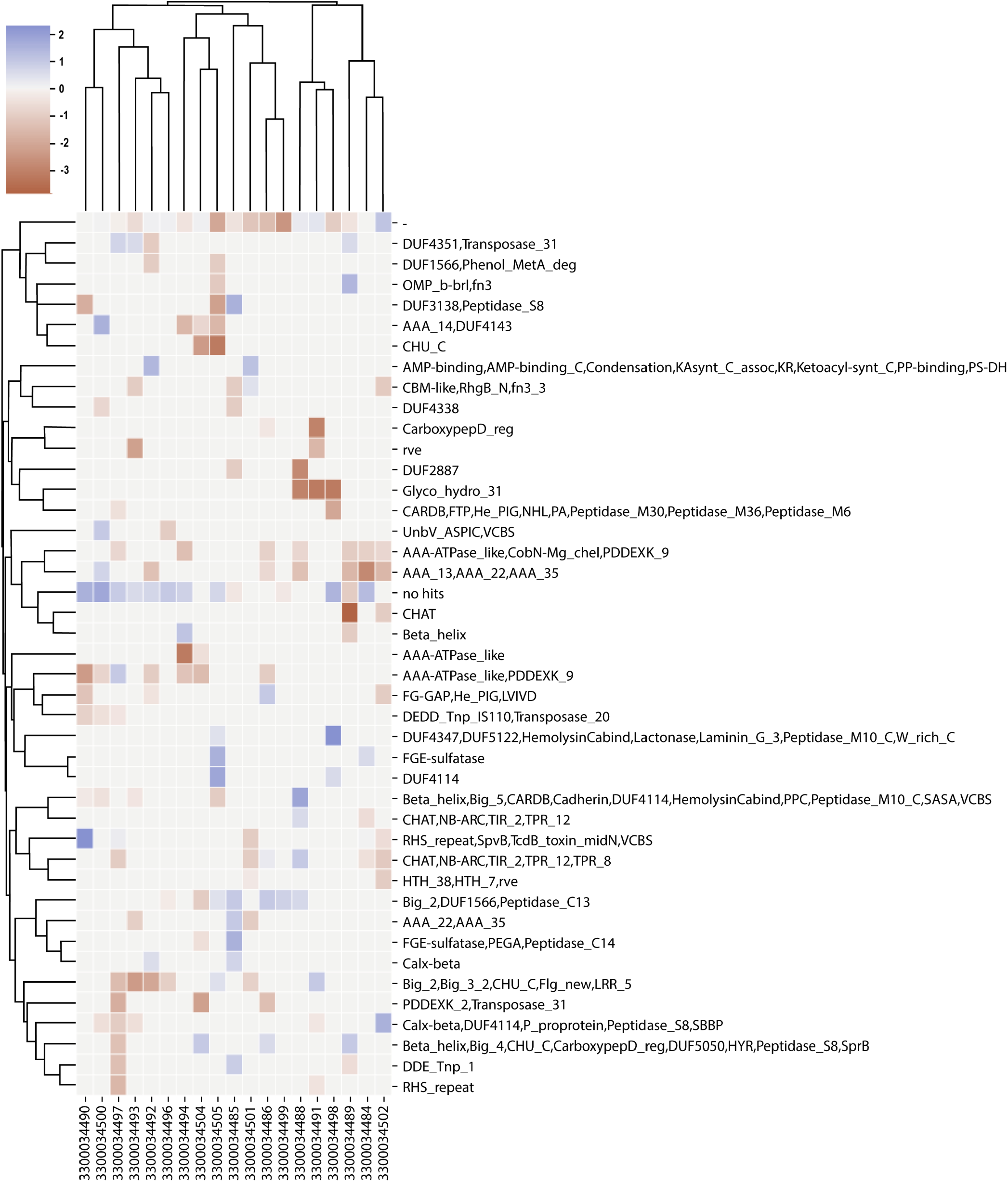
Heatmap and cluster analysis of pfams annotation of individual SNPs showing the log_2_ ratio of non-synonymous to synonymous substitutions (dN/dS) for the SNP differences contained within each SCM. The analysis suggested there was no positive selection of the protein-coding genes in which the SNPs were found.

**Fig. S7.**
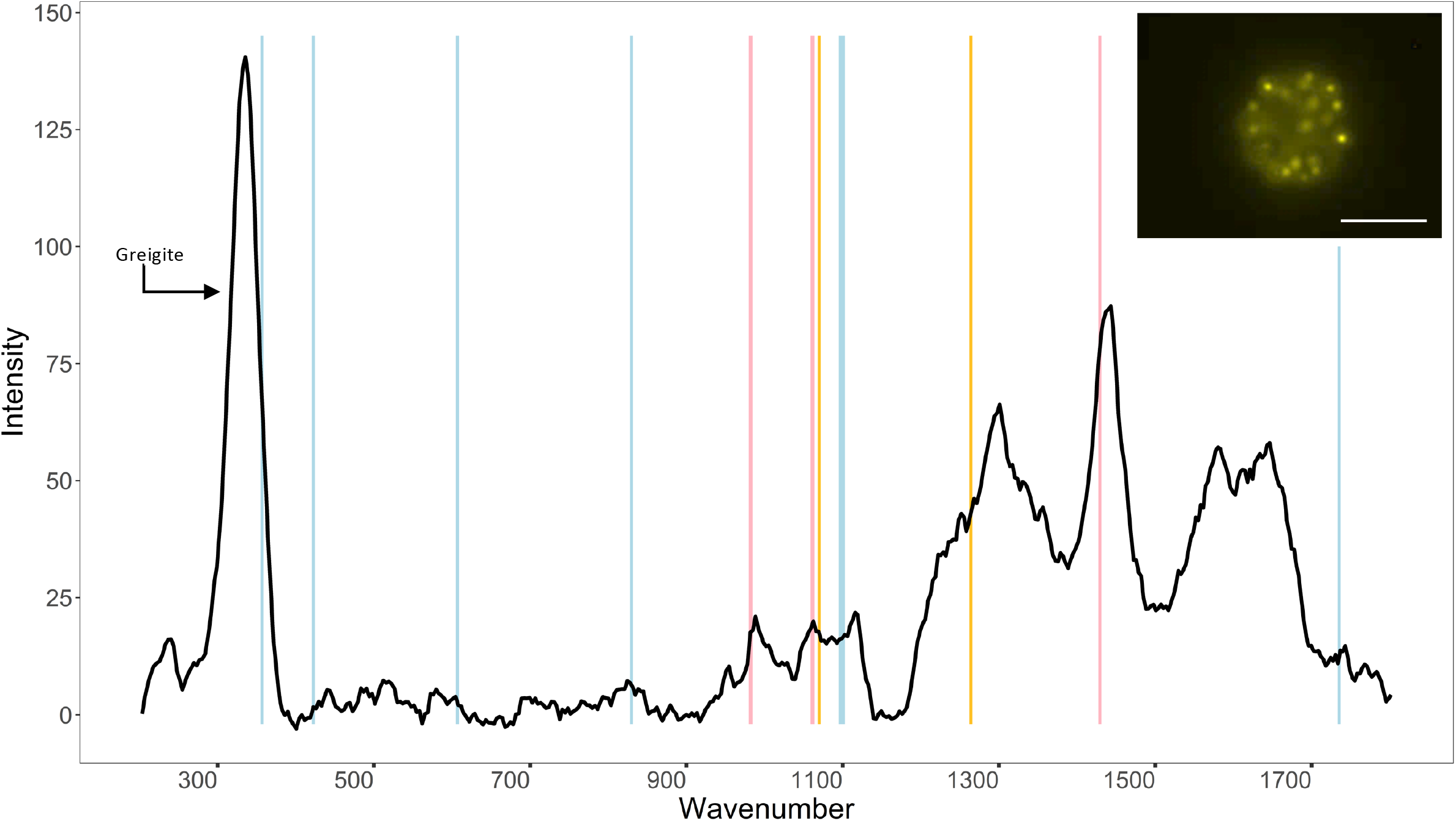
Representative Raman spectrum of a MMB using a 532 nm laser. Vertical lines show peaks corresponding to polyhydroxybutyrate (blue), triglycerides (gold), and exopolysaccharides (pink). Wavenumbers corresponding to peaks are listed in Table S6. The large peak at ∼335 cm^-1^ is assigned to the magnetosome crystal greigite, which has previously been shown for MMB from the same site (Schaible *et al*., 2022). Inset image shows a MMB consortium stained with Nile Red, indicating C-H rich droplets within cells. The contrast and brightness of the image has been increased for better visualization. Scale bar is 5 μm.

**Fig. S8.**
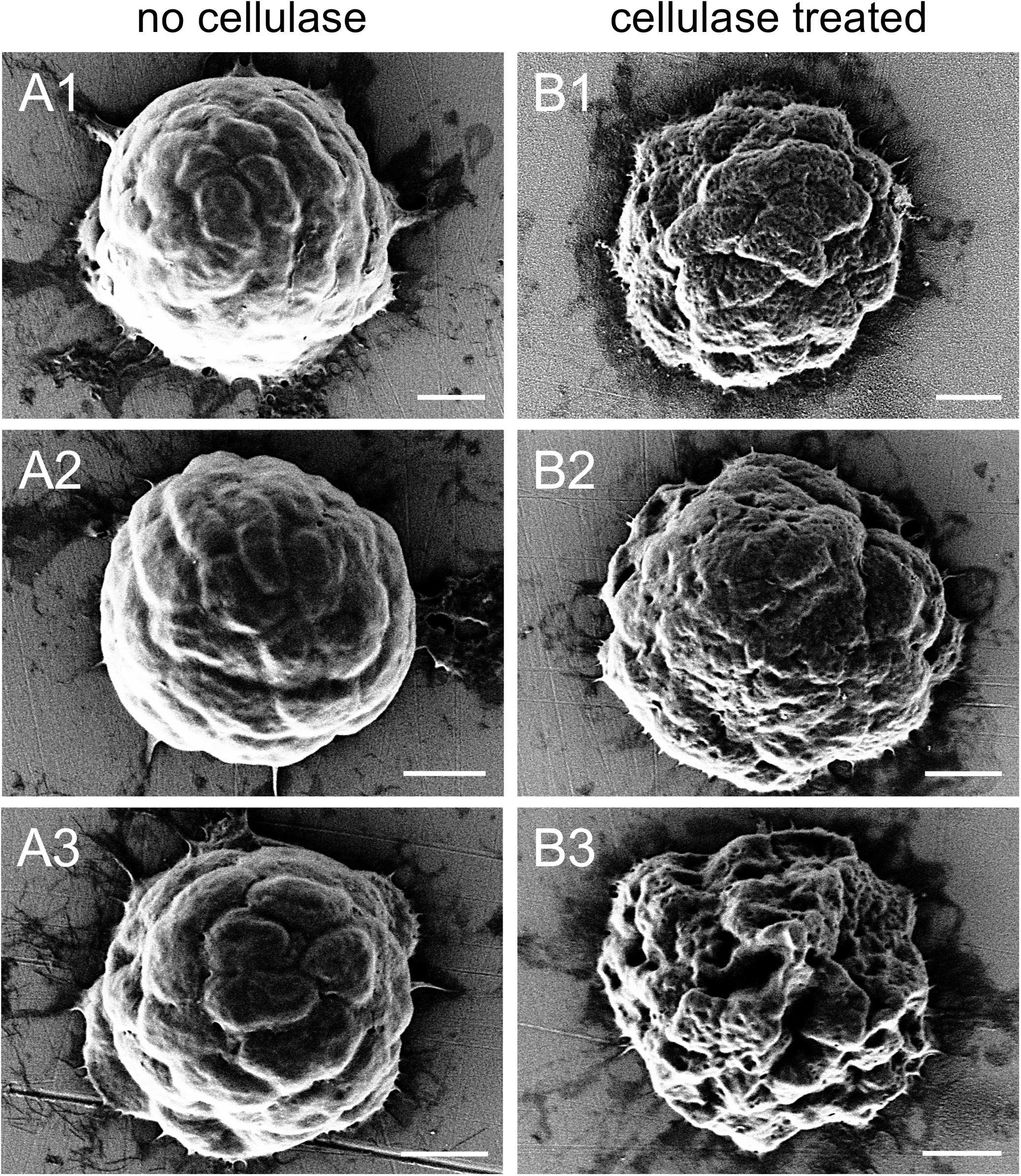
Cellulase treatment of MMB. (*A1-3*) Control sample of MMB incubated without cellulase. (B*1-3*) After treatment with cellulase the surface of MMB consortia was noticeably eroded as compared to the control. Both samples were incubated for 1 hr under otherwise identical conditions (pH, temperature, and osmolarity). All scale bars are 1 μm.

**Fig. S9.**
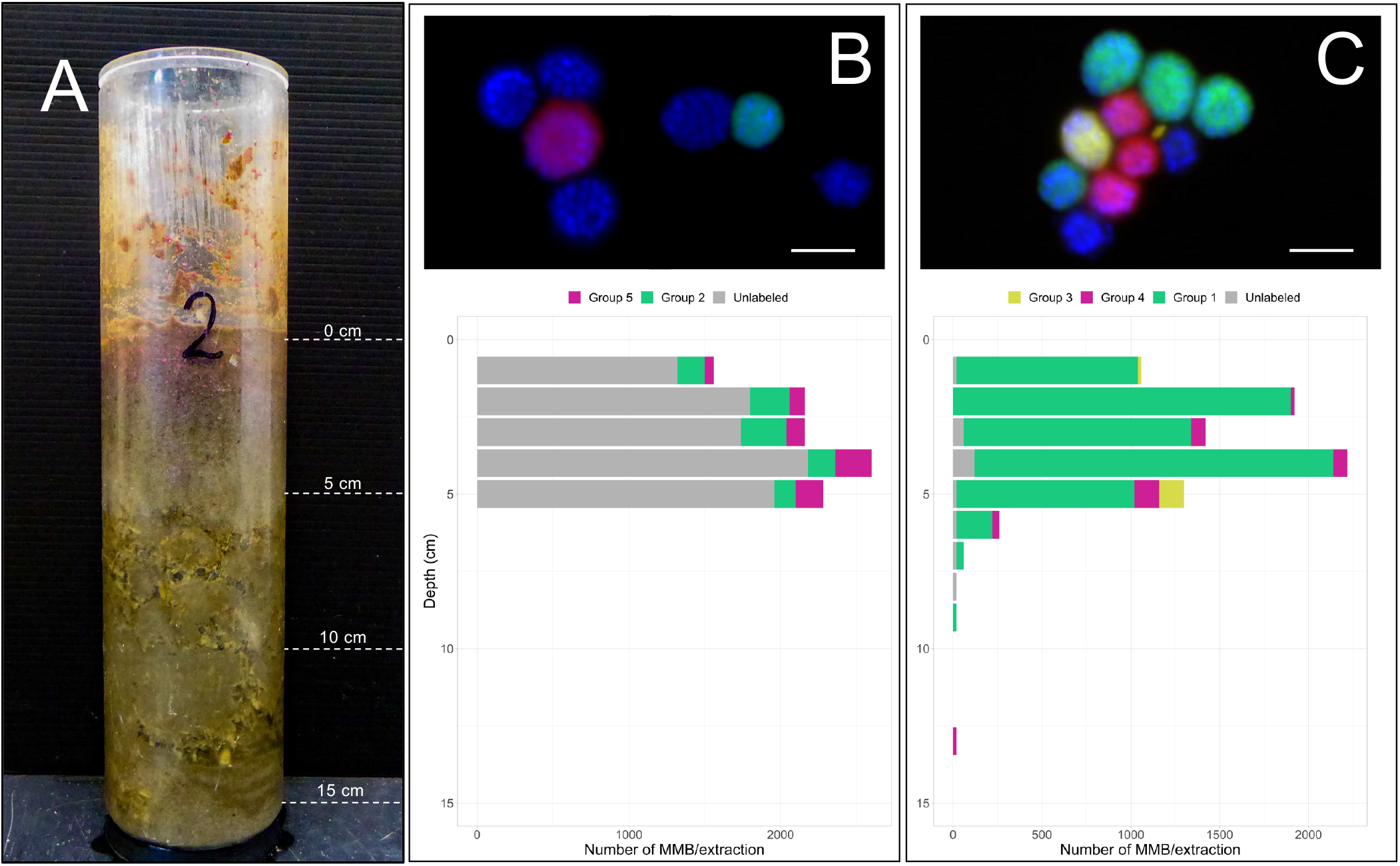
Fractional abundance of MMB groups by depth in LSSM. (*A*) Image of the 15 cm core taken from the West end of sampling site prior to being sectioned into 1 cm horizons from which MMB were enriched for quantification by FISH. (*B*) DOPE-FISH analysis of MMB Groups 2 (red) and 5 (green) shown in panel (*B*) and Groups 1 (green), 3 (yellow) and 4 (red) shown in panel (*C*). MMB not detected by the respective FISH probes are shown in the blue DAPI counterstain in the microscopy images. Scale bars are 5 µm. Bar plots show the abundance of each MMB group as determined by DOPE-FISH for each centimeter of the sediment core shown in panel (*A*). Unlabeled populations are MMB that were stained with DAPI but were not detected by the FISH probes used in the two separate experiments and are shown in gray. Consistent with results from SCM and previous 16S rRNA gene abundance studies (Simmons and Edwards 2007) in LSSM, Group 1 numerically dominate the MMB population. FISH probes used in this experiment are detailed in SI Appendix Table S9.

**Fig. S10.**
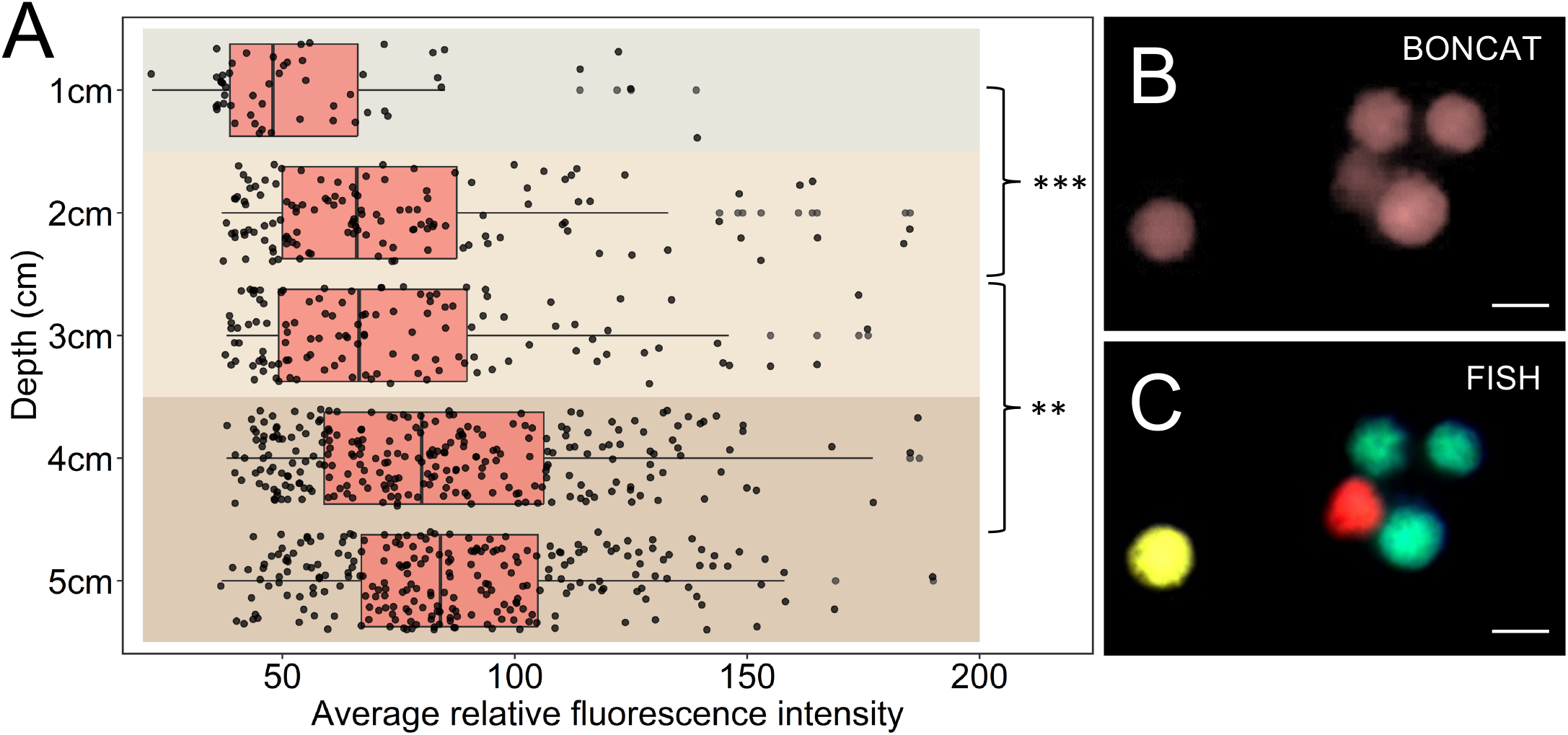
Anabolic activity of MMB inhabiting the top 6 cm of LSSM sediment as measured by BONCAT. *(A)* 1 cm sediment core horizons were incubated in the presence of the methionine analogue HPG and magnetically enriched MMB stained via azide-alkyne click chemistry with Alexa Fluor 405 to show relative activity of Group 1 MMB as a factor of depth in the sediment. The vertical line within each box shows the median and the whisker shows the range of the data. Dots represent individual MMB that were measured and analyzed using the software package Daime. Data points that were more than two standard deviations of the mean are shown as individual points past the whicker. The analysis showed that there is a statistically relevant difference in the activity of MMB from 1 cm depth to 2-3 cm and again from 2-3 cm to the 4-5 cm depth. (*B*) Exemplary epifluorescence microscopy image of click-stained MMB. (*C*) Overlay epifluorescence microscopy image of FISH-labeled MMB shown in panel *B*. Group 1 is shown in green, Group 3 in yellow, and Group 4 in red. All scale bars are 5 µm. All statistically differences are shown: ** = P < 3.9x10^-3^, *** = P < 3.5x10^-4^. FISH probes used in this experiment are detailed in SI Appendix Table S9.

**Fig. S11.**
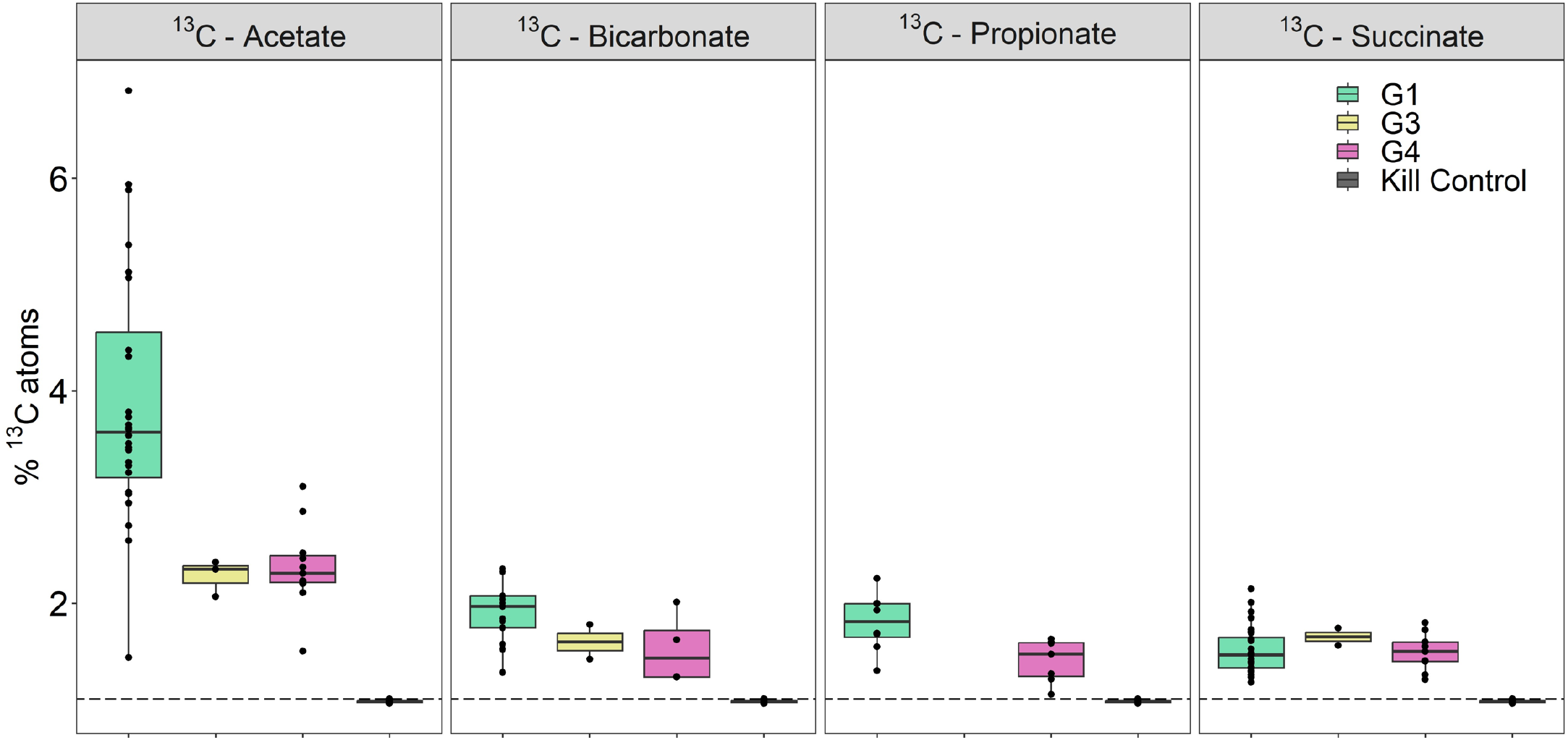
Comparison of ^13^C-labeled substrate incorporation by MMB Groups 1, 3, and 4 using NanoSIMS analysis of mass ratio ^13^C^12^C/^12^C_2_. The analysis shows that MMB in Group 1 anabolize acetate at a statistically greater rate than Groups 3 and 4 (p < 8.9x10^-3^). Group 1 also incorporated more bicarbonate than Group 4 (p < 2.4x10^-2^), although Group 4 only contained four samples to compare.

**Fig. S12.**
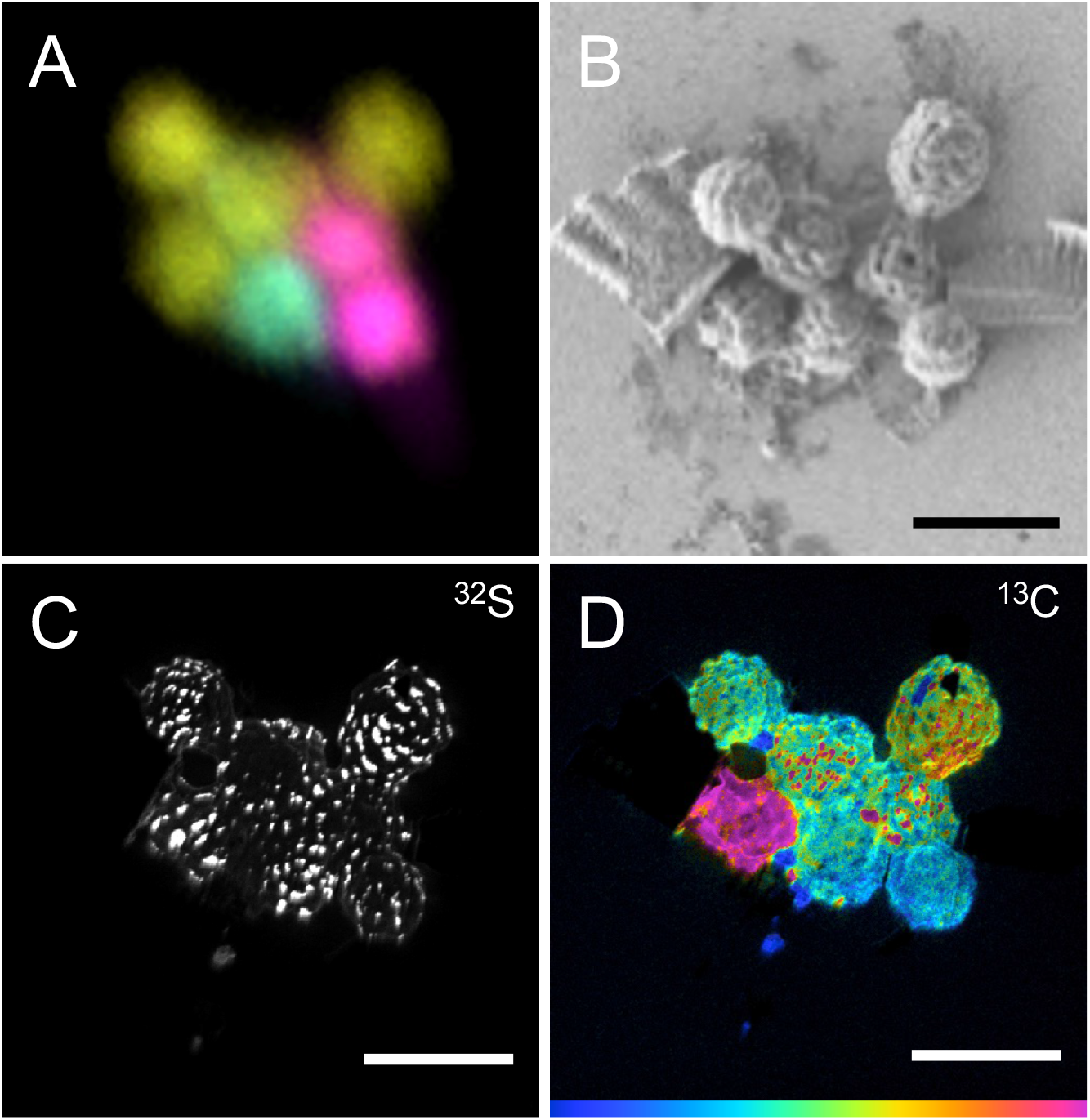
Correlative imaging of MMB to identify their *(A)* taxonomy (DOPE-FISH), *(B)* morphology (SEM), *(C)* distribution of sulfur (NanoSIMS, mass 32; a proxy for the presence of sulfur-containing magnetosomes) and *(D)* uptake of 1,2-^13^C_2_-labeled acetate (NanoSIMS, HSI image showing mass ratio ^13^C^12^C/^12^C_2_). Scale bars are 5 μm. Mass ratio color scale in *D* is 220-1000.

**Fig. S13.**
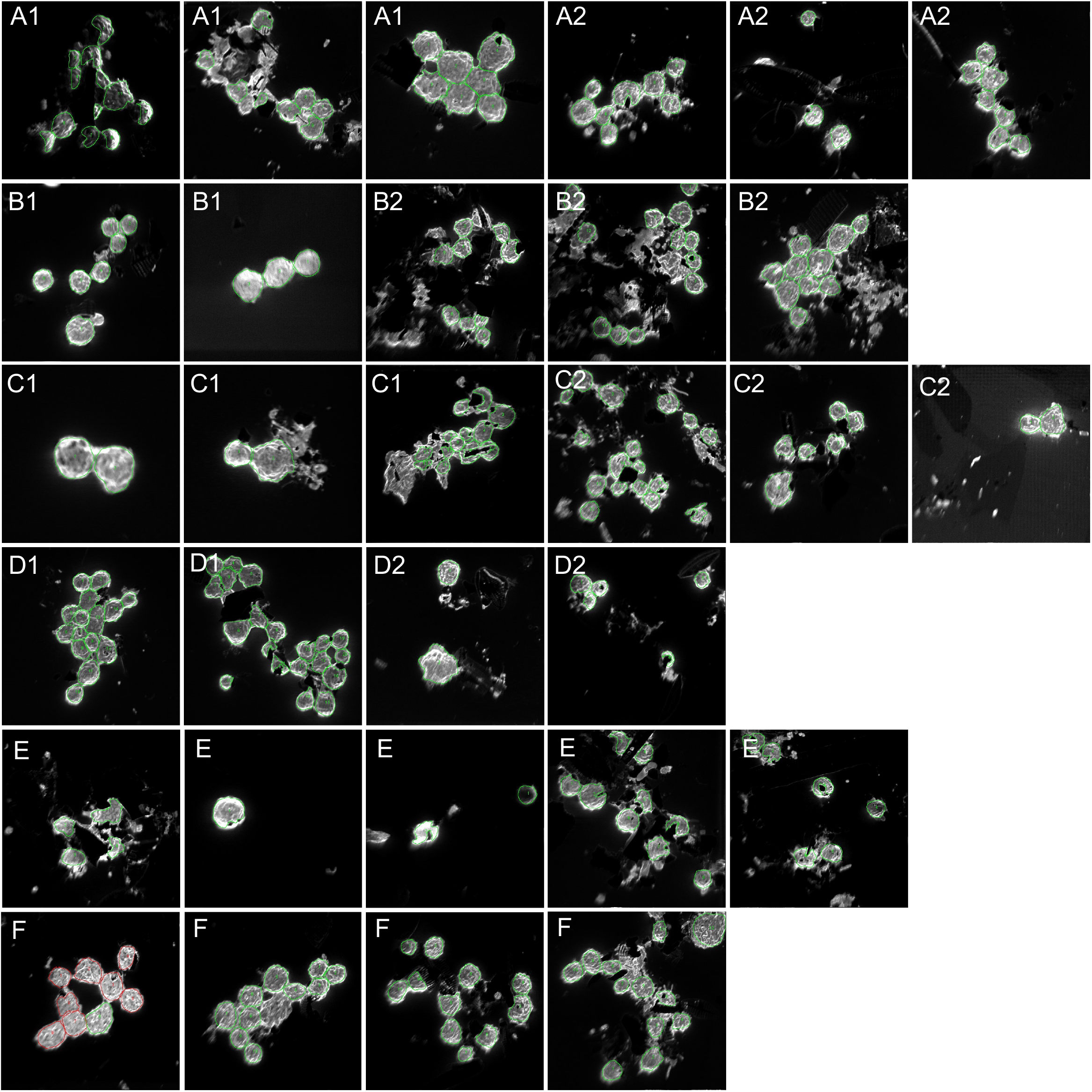
ROIs for NanoSIMS substrate analysis shown in Fig. 5 of main text. Because the *in situ* incubation incurred particles that were not of interest (*e.g*., diatoms and particulates), the ROIs were hand drawn around each MMB using the mass 26.00 (^12^C^14^N) channel as to avoid incorporation of exogenous material in the analysis. (*A1*) ^13^C-acetate, (*A2*) ^12^C-acetate, (*B1*) ^13^C-bicarbonate, (*B2*) ^12^C-bicarbonate, (*C1*) ^13^C-propionate, (*C2*) ^12^C-propionate, (*D1*) ^13^C-succinate, (*D2*) ^12^C-succinate, (*E*) ^13^C-acetate kill control, (*F*) negative control. ROIs are shown in green and red outlines.

**Fig. S14.**
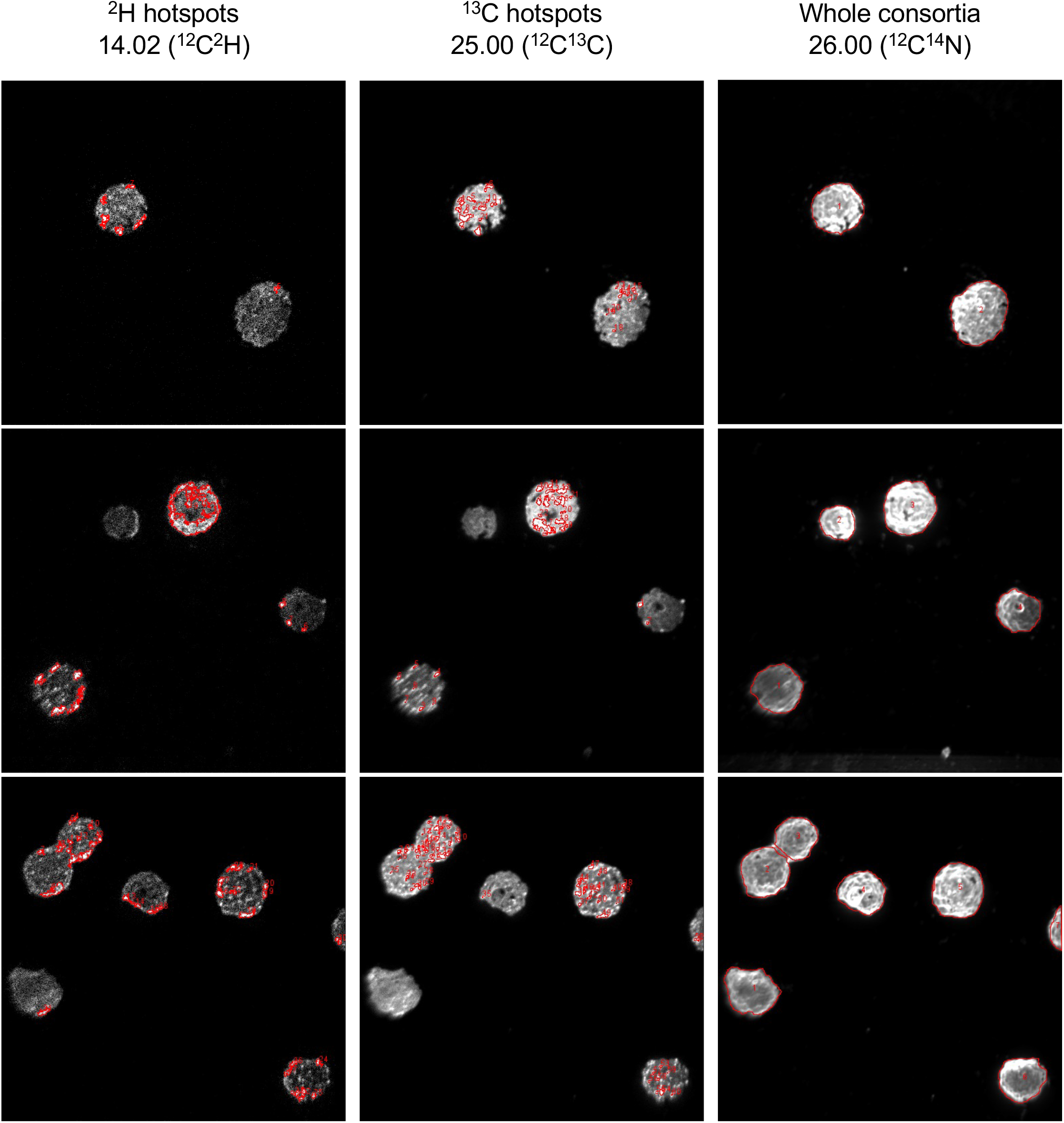
ROIs for NanoSIMS hotspot analysis shown in Fig. 6 of main text. As to avoid introducing bias into the selection of hotspot ROIs, thresholding in ImageJ was used to automatically select for ROIs, as outlined in the methods. The respective mass image was used for hotspot thresholding and ROI selection. ROIs for whole consortia were hand drawn. All ROIs are show in red outlines.

**Fig. S15.**
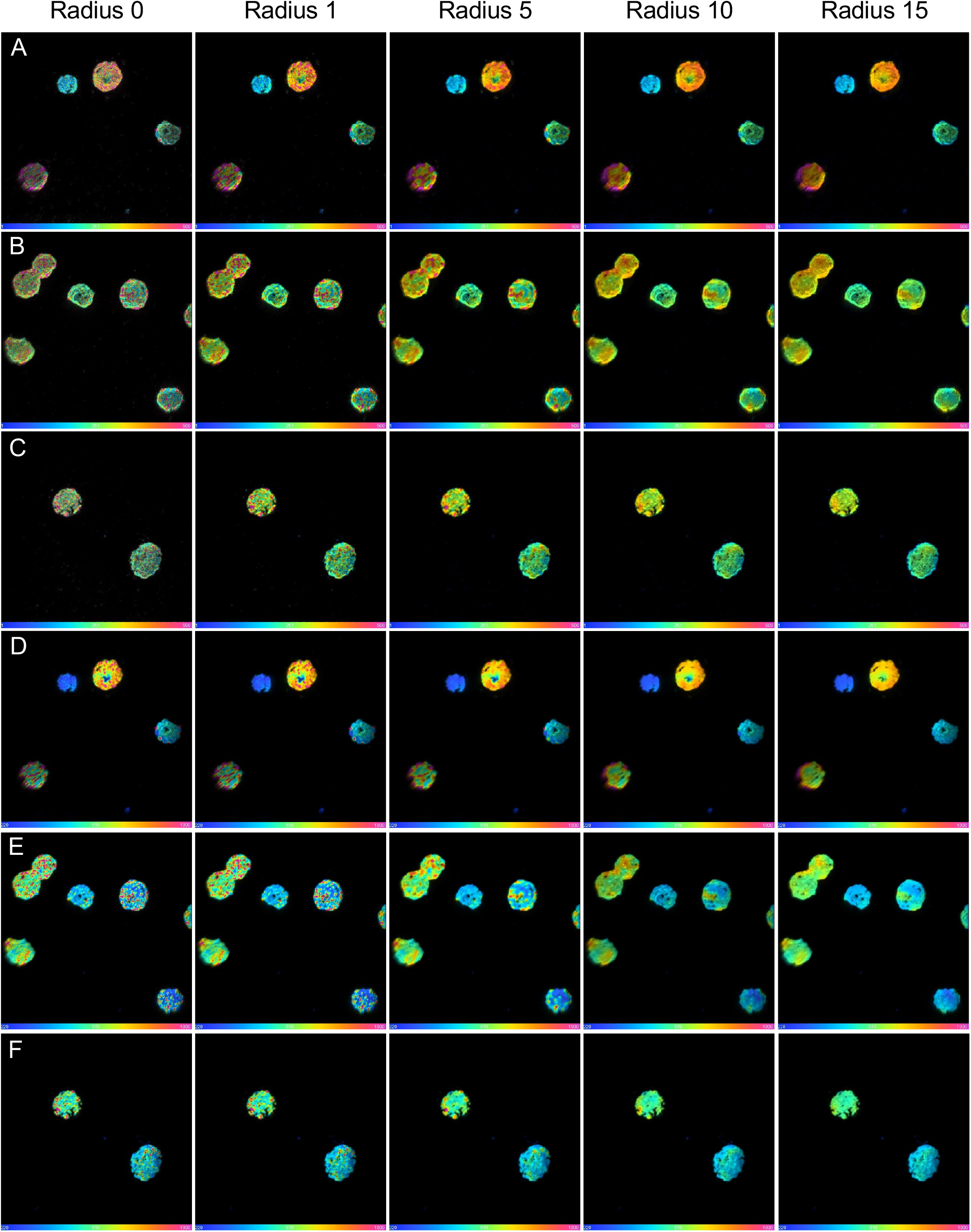
Median filter ratio radius effect on HSI NanoSIMS images of ^13^C and ^2^H hotspots *(A-C)* Mass ratio (^2^H^12^C/^1^H^12^C) of MMB labeled with deuterium oxide (^2^H_2_O). *(D-F)* Mass ratio (^13^C^12^C/^12^C_2_) of the same MMB shown in *A-C* but labeled with 1,2-^13^C_2_-labeled acetate. For these images, the median filter ratio radius was increased to show the effect of noise reduction and localization of isotope label within consortia. A higher filter radius reveals isolated areas of the respective isotope label within MMB, though for a radius > 5, the label is averaged over an area greater than the size of a single cell within the consortium, thus losing cellular resolution. Independent of the radius chosen, hot spots remain visible.

**Fig. S16.**
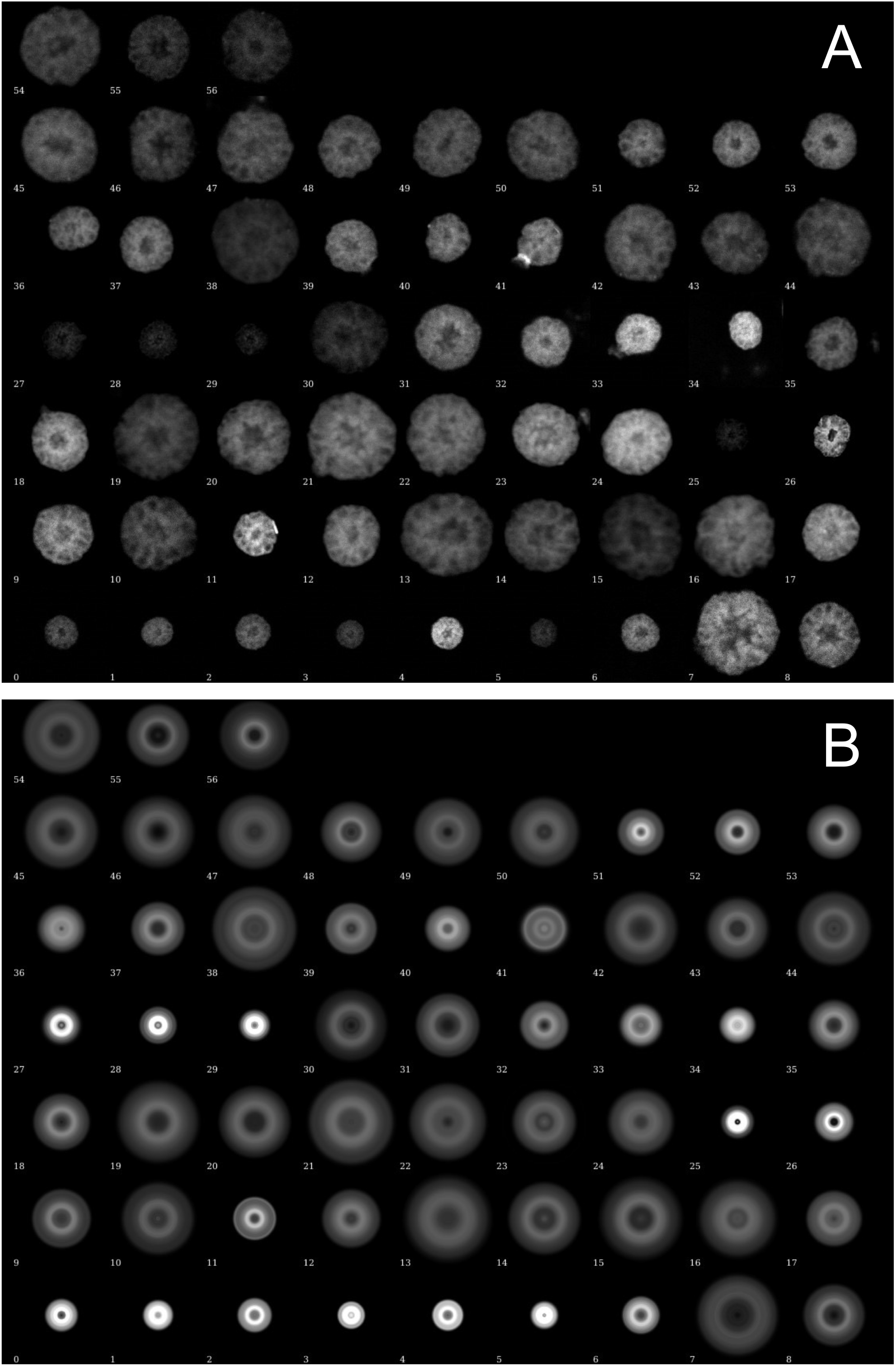
Anabolic activity within individual consortia. (*A*) Gray-scale images of individual MMB stained via azide-alkyne click chemistry with Alexa Fluor 488. (*B*) The same consortia shown in *A* that have been rotationally averaged in Eman2 software. The relative fluorescence intensity was standardized for all samples prior to analysis.

**Fig. S17.**
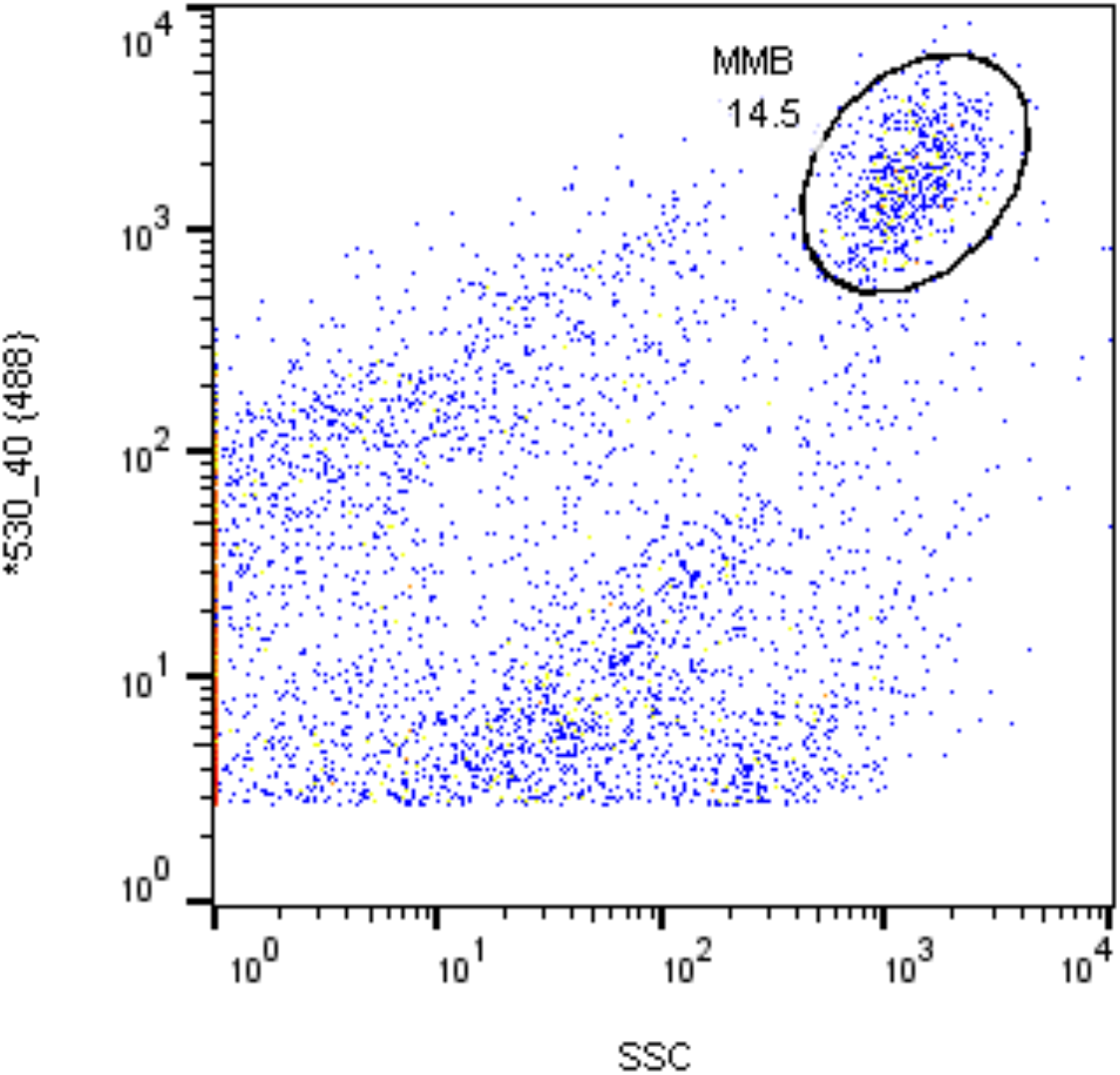
Fluorescence activated cell sorting of a magnetically enriched sample from tidal pond sediment stained with SYBR Green. A sorting gate, presumed to contain MMB consortia, was set around particles with a strong 488 nm signal and high side scatter (SSC), indicating a large cell size. Other particles likely are single cell magnetotactic bacteria or non-magnetotactic bacteria present in the pond water. MMB consortia were sorted into individual wells of a microtiter well plate and 22 MMB consortia were genome sequenced.

**Fig. S18.**
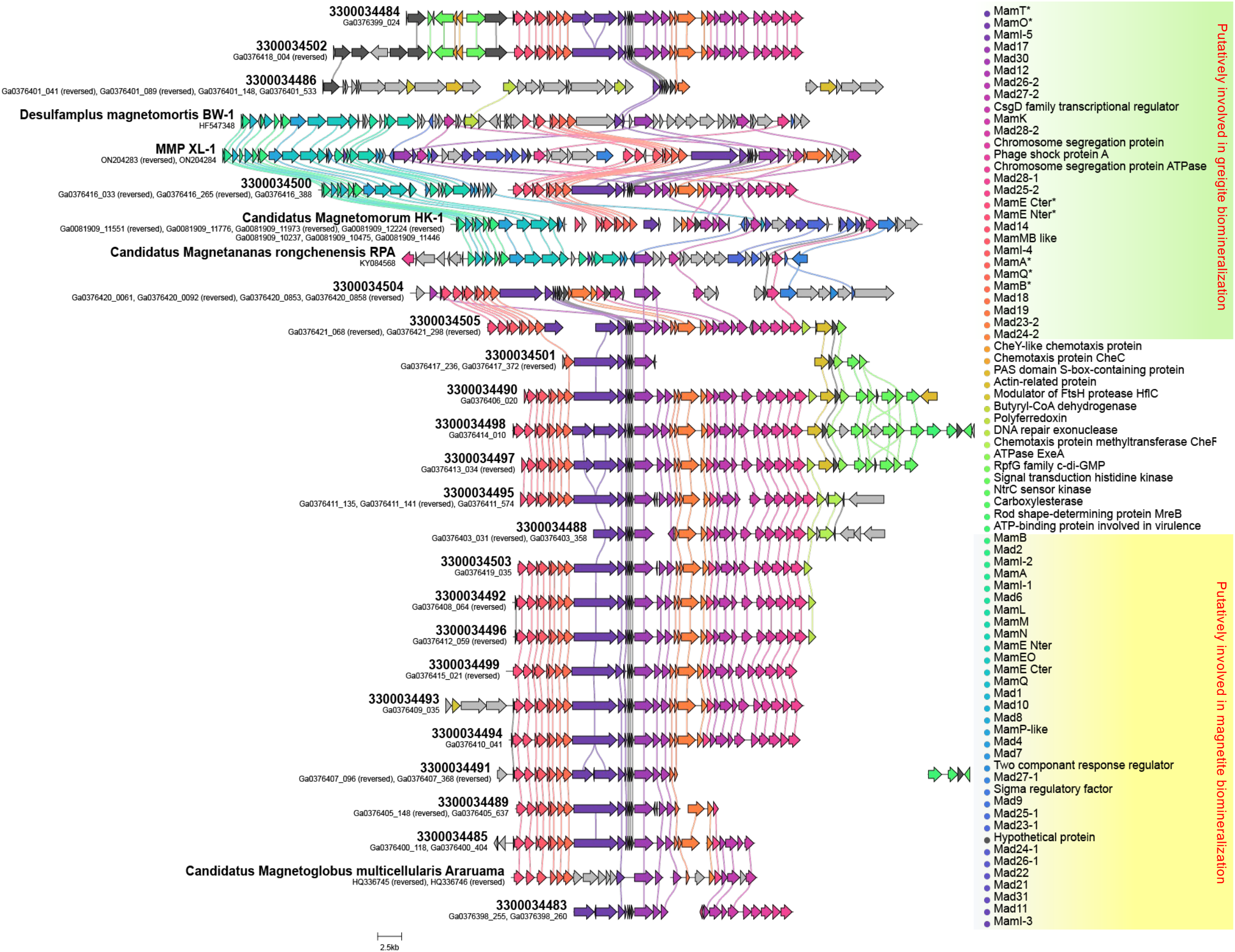
Gene synteny for scaffolds containing the magnetosome gene clusters compared. The corresponding annotations of colored genes are shown in the legend to the right.

